# Dissociating neural activity associated with the subjective phenomenology of monocular stereopsis: an EEG study

**DOI:** 10.1101/535203

**Authors:** Makoto Uji, Ines Jentzsch, James Redburn, Dhanraj Vishwanath

## Abstract

The subjective phenomenology associated with stereopsis, of solid tangible objects separated by a palpable negative space, is conventionally thought to be a by-product of the derivation of depth from binocular disparity. However, the same qualitative impression has been reported in the absence of disparity, e.g., when viewing pictorial images monocularly through an aperture. Here we aimed to explore if we could identify dissociable neural activity associated with the qualitative impression of stereopsis, in the absence of the processing of binocular disparities. We measured EEG activity while subjects viewed pictorial (non-stereoscopic) images of 2D and 3D geometric forms under four different viewing conditions (Binocular, Monocular, Binocular aperture, Monocular aperture). EEG activity was analysed by oscillatory source localization (beamformer technique) to examine power change in occipital and parietal regions across viewing and stimulus conditions in targeted frequency bands (alpha: 8-13Hz & gamma: 60-90Hz). We observed expected event-related gamma synchronization and alpha desynchronization in occipital cortex and predominant gamma synchronization in parietal cortex across viewing and stimulus conditions. However, only the viewing condition predicted to generate the strongest impression of stereopsis (monocular aperture) revealed significantly elevated gamma synchronization within the parietal cortex for the critical contrasts (3D vs. 2D form). These findings suggest dissociable neural processes specific to the qualitative impression of stereopsis as distinguished from disparity processing.

## Introduction

Viewing a pictorial image of a 3-dimensional (3D) scene produces a clear perception of depth relations and 3D object shape (Fig.1). However, the qualitative impression of depth and 3-dimensionality when viewing real scenes or stereoscopic images with both eyes is more compelling; there is a vivid sense of immersive negative space, tangible solid objects, and realness. This perceptual impression (stereopsis) is conventionally attributed to processing of binocular disparities to derive depth and 3D form (Ponce and Born, 2008; Westheimer, 2011; Wheatstone, 1838). However, there have been wide-ranging reports of the impression of stereopsis in the absence of binocular disparities, specifically when viewing a pictorial image with one eye through a reduction aperture (Ames, 1925; Koenderink, 1998; Michotte, 1991, 1948; Schlosberg, 1941; Vishwanath and Hibbard, 2013; da Vinci, cited in Wade et al., 2001; Wheatstone, 1838; see caption Figure 1), suggesting that it is not uniquely tied to processing of disparities. The phenomenological attributes associated with 3D visual experience under monocular aperture viewing by naive observers have been shown to be the same as those reported for stereoscopic viewing (Vishwanath and Hibbard, 2013). One conjecture is that the qualitative impression of stereopsis is not simply a by-product of disparity processing, but is associated with the conscious awareness of the efficacy of the visual representation for visually-guided manipulation (Michotte, 1948). Visually guided manual motor behaviour such as grasping and reaching requires precise representation of absolute (scaled) depth (Watt and Bradshaw, 2003). Since binocular disparity is the most effective cue for scaled depth representation at near distances, the above mentioned conjecture can explain why the impression of stereopsis is strongest in the presence of binocular disparity, while still accounting for its presence in other conditions where absolute depth could potentially be derived from a combination of cues other than disparity. Understanding the neural substrates that underlie the qualitative experience of stereopsis not only furthers our understanding of a fundamental aspect of our psychological visual experience of space (the tangibility, immersivity and realness associated with stereopsis) but may also in turn provide clues to the representation of 3D structure along the visual pathway.

**Figure 1.**
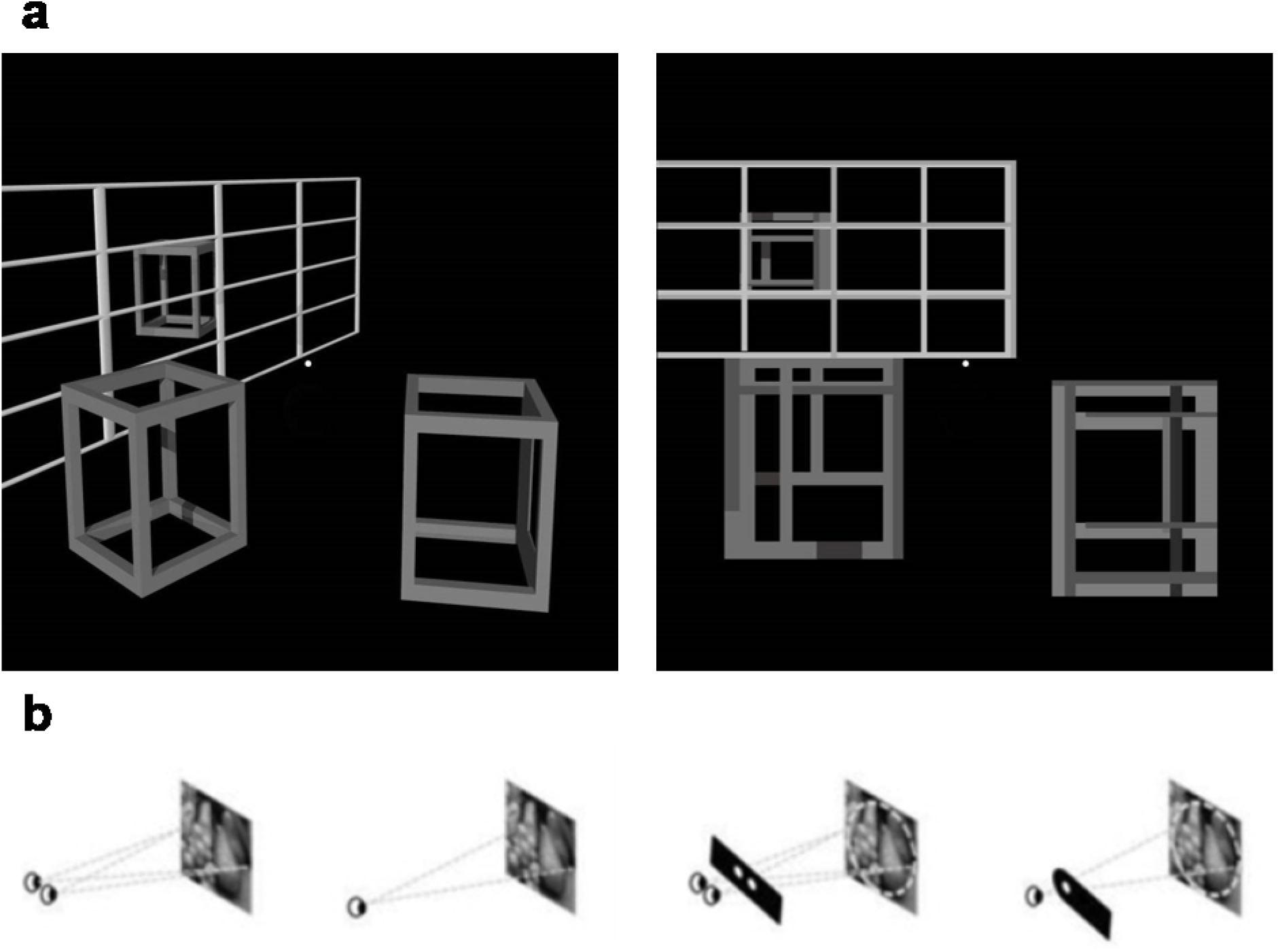
a) Examples of 3D (Left) and 2D (Right) geometric form images. b) Schematic representations of four different viewing conditions (Binocular, Monocular, Binocular Aperture, Monocular Aperture).

The primary focus of early studies in the neurophysiology of 3D depth perception had been on identifying and characterizing neural mechanisms of disparity processing (Barlow et al., 1967; Ohzawa et al., 1990; Poggio et al., 1985). Later work has focussed mainly on the neurophysiology of disparity derived depth and 3D structure (Backus et al., 2001; Ban and Welchman, 2015; Cumming, 2002; Durand et al., 2009; Georgieva et al., 2009; Goncalves et al., 2015; Neri et al., 2004; Preston et al., 2008; Taira et al., 2000; Tsao et al., 2003; Verhoef et al., 2011). While disparity and disparity derived depth studies had originally been based on single cell cat or monkey neurophysiology, more recently neuronal substrates underlying human 3D depth perception derived from binocular disparity has attracted much interest in neuroimaging studies, predominantly by functional Magnetic Resonance Imaging (fMRI) (see reviews Cumming and DeAngelis, 2001; Gonzalez and Perez, 1998; Orban, 2011; Parker, 2007; Sakata et al., 2005; Welchman, 2016; Welchman and Kourtzi, 2013).

Both monkey and human neuroimaging studies have provided important insight into the cortical landscape of neural responses to stimuli depicting 3D structure via binocular disparity. They demonstrate that 3D visual information is processed in both dorsal and ventral visual areas, but different information is processed in each stream (see a review, Goodale and Milner, 2018). It is suggested that the dorsal regions of the extrastriate cortex (V3A, V3B/KO, MT) and V7 are strongly engaged by disparity-defined depth information and involved in integrating signals to derive the 3D structure of viewed surfaces (Backus et al., 2001; Bridge and Parker, 2007; Cottereau et al., 2011; Durand et al., 2009; Goncalves et al., 2015; Minini et al., 2010; Naganuma et al., 2005; Preston et al., 2008), whereas the ventral regions of the extrastriate cortex (i.e. V3v, V4 and LOC, PPA) store representations of 3D scenes, object configurations and features (i.e. disparity-defined shape information) required for recognition and categorization (Bridge and Parker, 2007; Chandrasekaran et al., 2007; Gilaie-Dotan et al., 2002; Kourtzi and Kanwisher, 2001; Neri et al., 2004).

The majority of the studies examining binocular stereopsis both in monkey and human neurophysiology have used random dot stereograms (RDSs: Julesz, 1971) representing disparity-defined depth/object/shape when viewed stereoscopically (Human: Backus et al., 2001; Ban and Welchman, 2015; Goncalves et al., 2015; Neri et al., 2004; Preston et al., 2008; Monkey: Cumming, 2002; Taira et al., 2000; Tsao et al., 2003; Verhoef et al., 2011). Since RDSs produce an impression of depth and stereopsis in the presence of binocular disparities, using this paradigm (or random line stereograms, e.g., Durand et al., 2007) makes it challenging to distinguish between neural processes underlying disparity processing, the derivation of depth/3D shape and those that underlie the qualitative impression of stereopsis. Importantly, using RDS stimuli precludes distinguishing between conditions that generate depth perception without the impression of stereopsis (pictorial depth, Fig.1) and those that produce depth perception along with the qualitative impression of stereopsis (binocular stereopsis or monocular stereopsis, Fig.1).

The suggestion that the qualitative impression of stereopsis might involve distinct processes leading to conscious awareness of the capacity for precision visually guided manipulation (e.g. Michotte, 1948) implicates higher-level cortical areas beyond the extrastriate cortex, particularly the parietal cortex. The dorsal visual pathway extends from the primary visual cortex in the occipital lobe to the parietal lobe, which is important in providing sensory input for movements (see reviews Anzai and DeAngelis, 2010; Freud et al., 2016; Gallivan and Culham, 2015; Goodale and Milner, 2018; Tunik et al., 2007). Additionally, the parietal cortex is thought to support an extensive range of sensory and cognitive functions including spatial representation, multimodal integration, attentional control, motor planning, and working memory (see reviews, Behrmann et al., 2004; Buneo and Andersen, 2006; Culham and Kanwisher, 2001; Grefkes and Fink, 2005; Hubbard et al., 2005; Kastner and Ungerleider, 2000; Merriam and Colby, 2005; Mesulam et al., 2005; Orban et al., 2006; Wagner et al., 2005; Yantis and Serences, 2003). Most importantly, accumulating evidence of human fMRI studies demonstrates that the human parietal cortex processes visual information in a form that supports the ability to manipulate objects both physically (Binkofski et al., 1998; Culham et al., 2003; Freud et al., 2018; Shikata et al., 2003; Snow et al., 2011) and mentally (Gauthier et al., 2002; Shikata et al., 2003).

Non-human primate fMRI studies have consistently shown strong activity in the parietal cortex during 3D perception in the presence disparities (Durand et al., 2007; Joly et al., 2009; Rosenberg et al., 2013; Rosenberg and Angelaki, 2014; Taira et al., 2000; Tsao et al., 2003; Tsutsui et al., 2002; Van Dromme et al., 2016, 2015; Verhoef et al., 2015). However, only a few fMRI studies (e.g., Chandrasekaran et al., 2007; Durand et al., 2009; Georgieva et al., 2009; Minini et al., 2010; Tsao et al., 2003) have examined the processing of 3D shape from binocular disparity in parietal cortex in humans. Georgieva et al. (2009) found several intraparietal sulcus (IPS) regions (i.e. the dorsal IPS anterior (DIPSA), the dorsal IPS medial (DIPSM), the parieto-occipital IPS (POIPS) and ventral IPS (VIPS) regions) to be activated in the presence of binocular disparity defined depth structure. Interestingly, these regions had been previously shown to be involved in the perception of 3D shape from motion (Murray et al., 2003; Orban et al., 1999; Vanduffel et al., 2002), from texture (Georgieva et al., 2008; Shikata et al., 2001; Tsutsui et al., 2002) and shading (Taira et al., 2001). Taken together, these findings show that significant processing for the derivation of 3D structure and its subsequent use for manual action occurs in the parietal cortex.

However, there have been few studies that have specifically aimed to determine at what stage of the dorsal visual stream, information from depth cues (e.g. disparity) for deriving 3D structure becomes converted into a format for the guidance of action; for example, the conversion of relative depth information in the cues into absolute depth values (Watt and Bradshaw, 2003). One possibility is that the processing involved in the generation of absolute depth values required for manual action also underlie the qualitative impression of stereopsis. For example, discrimination of depth on the basis of reaching actions has been shown to be possible under monocular aperture viewing (condition that elicits the phenomenology of stereopsis) but not normal binocular viewing of pictorial images (Volcic et al., 2014). Understanding the neural substrates that underlie the qualitative experience of stereopsis may provide indirect clues to the representation of 3D structure. A critical step in this understanding is to distinguish areas in the visual stream that are activated when 3D structure is perceived in the presence of stereopsis but not in the absence of stereopsis (i.e., the perception of pictorial depth; see Figure 1).

To this end, we sought to distinguish between neural processing for cases that generate only the perception of pictorial depth without stereopsis (binocular viewing of pictorial images; Fig 1) or generate an impression of depth with the quality of stereopsis (monocular aperture viewing; Fig.1), while excluding the potentially confounding role of binocular disparities. In order to do this we aimed to examine changes in oscillatory activity in different frequency bands (i.e. delta 1-3Hz; theta 4-7Hz; alpha 8-13Hz; beta 14-30 Hz; gamma 30-100 Hz) to non-stereoscopic images of either 2D or 3D pictorial content under different viewing conditions (monocular, binocular, monocular aperture, binocular aperture).

Recent studies have linked neural synchronization in distinct frequency bands with a range of cognitive and sensory functions (Buschman and Miller, 2007; Buzsaki and Draguhn, 2004; Colgin et al., 2009; Fries, 2009; Jensen et al., 2014; Klimesch, 1999; Pfurtscheller et al., 1996; Pfurtscheller and Lopes da Silva, 1999; Singer and Gray, 1995). It is well known that low-frequency oscillations (i.e. alpha or beta) are suppressed by sensory stimulation (Bauer et al., 2006; Pfurtscheller and Lopes da Silva, 1999), whereas high-frequency oscillations (i.e. gamma) are enhanced (Donner and Siegel, 2011; Hoogenboom et al., 2006).

Lower frequency activity, typically alpha activity, has been suggested to reflect functional inhibition of neural processes by suppressing irrelevant incoming sensory signals (Bauer et al., 2014, 2006; Haegens et al., 2011; Jensen and Mazaheri, 2010; Klimesch et al., 2007; Pfurtscheller and Lopes da Silva, 1999; Spaak et al., 2012; Thut and Miniussi, 2009), while beta activity is thought to have different functional characteristics related to the processing of relevant stimuli in large-scale networks (see reviews Fries, 2015; Kilavik et al., 2013; Siegel et al., 2012; Wang, 2010). On the other hand, increase in the power of neural oscillations in the gamma frequency band is a key signature of information processing in cortical neural network subserving fundamental operations of cortical computation (Donner and Siegel, 2011; Fries, 2009; Muthukumaraswamy and Singh, 2013). Gamma-band synchronization has been observed in humans using non-invasive imaging methods during visual (Hoogenboom et al., 2006; Muthukumaraswamy and Singh, 2013), somatosensory (Bauer et al., 2006) and auditory (Pantev et al., 1991; Schadow et al., 2009) stimulation. It is also known to be involved in higher cognitive functions such as memory processes (Fell et al., 2001; Howard et al., 2003) and motor control (Brown et al., 1998; Cheyne et al., 2008; Crone et al., 1998; Darvas et al., 2010; Gaetz et al., 2010; Muthukumaraswamy, 2010; Schoffelen et al., 2005). In addition, specifically with respect to visual processing, recent studies in the primate (Bastos et al., 2015; Mejias et al., 2016; Ni et al., 2016; Richter et al., 2017; van Kerkoerle et al., 2014) and human (Michalareas et al., 2016) have demonstrated that gamma oscillation is related with feedforward (bottom-up) processes, whereas alpha/beta activities carried feedback (top-down) signals. The directionality of neural communications has started to explain the role of distinct neural oscillations in terms of feedforward (bottom-up) (ascending hierarchical levels) and feedback (top-down) (descending hierarchical levels) signal processing (Bastos et al., 2012; Buschman and Miller, 2007; Engel et al., 2001; Fries, 2015; Scheeringa et al., 2016; Wang, 2010).

This accumulating evidence consistently reveals that high- and low-frequency oscillations show different responses to afferent input reflecting distinct neural processes. We therefore focussed our analysis on broad power changes in high and low frequency domain in order to detect a dissociable neural signature of the experience of the stereopsis. Methodologically, we localized EEG oscillatory sources (beamformer techniques) in the visual and parietal cortex, and conducted time-frequency analysis to examine power change in alpha (8-13Hz) and gamma (60-90Hz) frequency in both regions. We anticipated that the dissociable neural activity associated with depth perception would be most likely identified in gamma frequency band based on previous literature that 3D processing is largely a stimulus driven bottom-up process (Cumming and DeAngelis, 2001; Gonzalez and Perez, 1998; Orban, 2011; Parker, 2007; Welchman, 2016; Welchman and Kourtzi, 2013).

## Methods

### Participant

14 right-handed subjects (4 males, 10 females, age = 24.5±2.8), who were naive to the purposes of the experiment, took part in the study. All participants gave informed written consent. The experiment was reviewed and approved by the University Teaching and Research Ethics Committee in the School of Psychology and Neuroscience at the University of St Andrews. All procedures were conducted in accordance with the ethical guidelines of the University of St Andrews and the 1964 Declaration of Helsinki. Each participant participated in a single session lasting approximately two hours, including EEG setup.

First, all participants were screened for their visual acuity. They were asked to read a Western Optical Standard Reading Card (Western Ophthalmics, Lynnwood, WA, USA) with either normal or corrected-to-normal vision if necessary, and visual acuity was deemed sufficient if participants could clearly read text that subtended a visual angle of five inches at 20 inches distance. Second, participants were screened for their binocular stereo-acuity. They completed a Randot Stereotest (Stereo Optical Co., Chicago, IL, USA), and binocular stereo-acuity was deemed sufficient if participants could identify geometric forms at 500 and 250 seconds of arc from 16 inches, and could complete the graded circle test to at least level eight which is 30 seconds of arc from 16 inches. One participant was excluded due to the inability to identify the geometric forms and was not tested further. The remaining participants were then assessed for their ability to experience monocular stereopsis when viewing a pictorial image though a reduction aperture (see Vishwanath and Hibbard, 2013). They successively viewed two natural photographs of 3D leaves or ferns (1920 x 1080 pixels) on an LCD monitor under binocular and monocular-aperture viewing. Participants were asked to indicate whether they perceived any difference in depth impression between the two viewing conditions, and if do they were asked to indicate in which one they perceived a more compelling impression of depth. Then, using a Likert-type questionnaire (Vishwanath and Hibbard, 2013) they were asked to agree or disagree with a set of statements regarding the perceptual differences they perceived. The ability to perceive monocular stereopsis was deemed sufficient if participants reported better qualitative experience under monocular aperture viewing and answered in the affirmative to the items consistent with phenomenal experience of stereopsis but not sham items in the likert questionnaire (Vishwanath and Hibbard, 2013). One participant, despite possessing normal binocular stereo-acuity, reported no difference in depth impression between the two viewing conditions, was excluded and not tested further. The inability to perceive monocular stereopsis of a single participant in this sample size (N=14) was expected and consistent with the previous research, which found that approximately 10% of 60 participants could not perceive monocular stereopsis (Vishwanath and Hibbard, 2013). After the visual screening, twelve participants were tested in the main experiment.

### Stimuli

Stimuli consisted of a set of 16 3D and 16 2D geometric form greyscale images (see Figure 1a) for a total 32 images. 3D images were produced using the raytracing and modelling software POVray Version 3.7. Each 3D image had a black background and contained three hollow cubes – two in the foreground and one in the background – and a lattice frame that ran from the side of the image and ended approximately in the centre of the image. Variations between images were in the size, orientation, and position of the cubes; the orientation and number of horizontal and vertical bars in the lattice; and the position of the virtual light source. The 2D form images were constructed using PowerPoint 2013 such that the 2D shapes matched as closely as possible the size, location, spatial distribution, greyscale lightness values of component elements in the 3D version, but without containing any cues to pictorial depth, including interposition.

### Display Apparatus

The stimuli were presented on a 17-inch CRT monitor (Sony CPD-200ES) operating with a resolution of 800 x 600 pixels and a refresh rate of 75Hz, controlled by a Pentium PC. Participants’ heads were stabilized using a chin rest at a viewing distance of 80cm. They wore a pair of specially designed black plastic spectacles into which card slides were inserted to create four different viewing conditions: binocular (B), monocular (M), binocular-aperture (BA), and monocular-aperture (MA) (see Figure 1b). Participants viewed with their dominant eye in any monocular conditions. The temples of the spectacles were designed to block the peripheral visual field. In the aperture conditions, participants viewed the stimuli through oval apertures (4 x 2mm), the centre of which were located 6mm above the nose bridge of the spectacles, to align approximately with the centre of the pupil, and approximately 15mm in front of participants’ eyes, occluding the monitor frame. Participants individually adjusted the aperture slides horizontally until a fixation cross in the centre of the monitor was centred in the apertures, and they could not see the boundaries of the image. In the conditions without the aperture, the monitor frame was visible. Stimulus display was programmed with Experimental Run Time System (ERTS) software (Beringer, 1992).

### Data acquisition

EEG data was acquired using a BIOSEMI Active-Two amplifier with 70 Ag/AgCl electrodes. Four electrodes of them were placed at the outer canthi of and below each eye to record electrooculography (EOG). The electrode layout followed the extended international 10-20 system. The Common Mode Sense (CMS) and driven right leg (DRL) electrodes were utilized as an active noise cancellation system (http://www.biosemi.com/faq/cms&drl.htm for details). Data were recorded at a sampling rate of 256 Hz.

### Experimental Paradigm

#### Visual detection task

The experimental paradigm and the stimuli are illustrated in Figure 2. In order to maintain a uniform state of visual fixation and attention, the display of the images was combined with a visual detection task at fixation. Each trial started with the presentation of a white fixation dot (2 x 2mm) at 5mm below the centre of the monitor. After 1500ms, a geometrical form image (3D or 2D) appeared (the white fixation dot remained visible). 800ms after the image onset, the white fixation dot changed colour to either orange or green. The subjects’ task was to indicate the colour the fixation dot had changed to. The participants were instructed to respond with their index finger of each hand using two ERTS response keys, 15cm apart horizontally. Immediately following subjects’ response or after 4000ms if participants had not responded, the image of geometrical shapes disappeared, and the fixation dot changed colour back to white, and after 1500ms, the next image appeared. Responses faster than 150ms or slower than 1500ms relative to the colour change onset were excluded from analysis. The subjects were instructed to respond both quickly and accurately, and to fixate on a fixation dot, remaining as still as possible throughout the whole experiment. Assignment of the orange and green colour change to left and right button was counterbalanced across participants to control for effects of the response key positioning.

**Figure 2.**
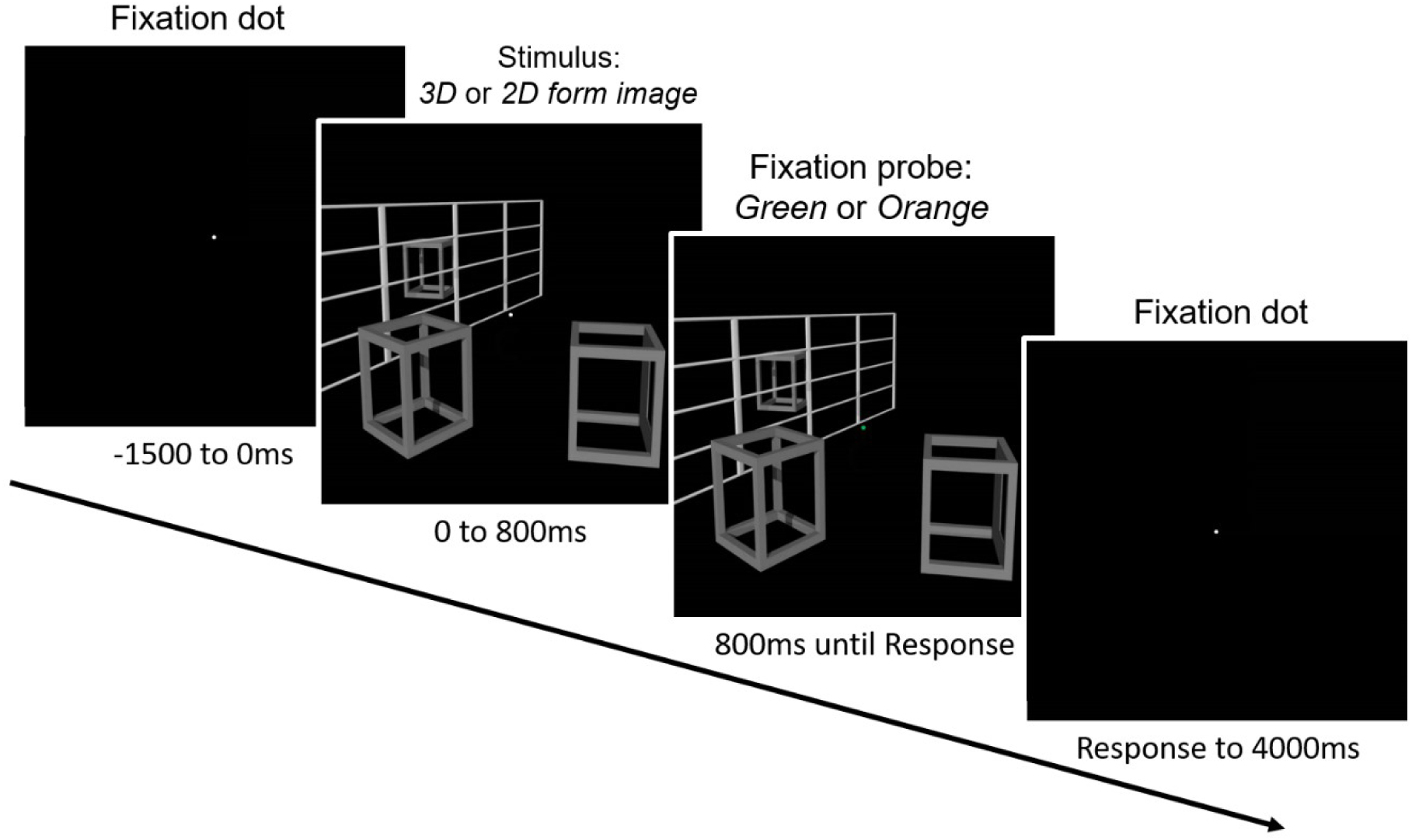
Timeline of stimulus presentation and fixation detection task (see text for details).

There was an initial practice block of eight trials followed by eight experimental blocks of 64 trials each. For each block, 32 geometrical form images (sixteen 3D and sixteen 2D images) were presented twice each in random order. For each of the four viewing conditions (M, B, MA, BA), there were two consecutive blocks, resulting in 64 trials for each 3D and 2D form images under each viewing condition.

The order of viewing conditions was randomised across participants using a Latin square design. At the end of each block, visual feedback was given about their mean reaction time (in ms to one decimal place, including the probe visual cue period of 800ms) and the number of errors they had made, as incentives to concentrate on the task).

#### Stereopsis impression rating task

After the completing the eight EEG recording blocks, participants remained in the identical position for a final visual depth impression rating task. Participants were asked to rate subjective strength of the stereopsis impression when viewing 3D form images under each viewing condition. To operationalise subjective impression, firstly participants were asked to self-define a rating scale of 1 to 5 (see Vishwanath and Hibbard, 2013). On this scale, 1 was defined as the strength of stereopsis under binocular viewing of the 3D form image (no stereopsis), whereas 5 was defined as the strength of stereopsis under monocular aperture viewing (paradigmatic monocular stereopsis). Participants were allowed to view the image between these two viewing conditions as many times as they wished until they were confident that they had internalised the difference in subjective impression of depth at the two ends of the scale. Participants then viewed the same image under binocular aperture and monocular viewing conditions, and provided verbal ratings of the strength of stereopsis impression in relation to the self-defined scale of 1 to 5. This procedure was repeated for each four 3D images.

### Data analysis

#### Behavioural data

Response Accuracy (% Correct) and Reaction Time (ms) were measured during the visual detection task. The RT was the time between the onset of the fixation colour change and key press (only correct trials were included in the RT analysis). Data were averaged across trials for each condition (4 Viewing x 2 Image dimension) for each subject. Both dependent variables were submitted to separate 2 type of vision (Monocular, Binocular) x 2 aperture condition (Aperture, Non-aperture) x 2 image dimension (2D form, 3D form) repeated-measures ANOVA. In the event of a significant interaction effect, the Bonferroni method was used to adjust the p value of post hoc pairwise comparisons.

#### EEG data

We focused our analysis a-priori based on the previous literature, which suggested that depth representation processes would be a bottom-up feedforward processing occurring within the visual cortex and/or parietal cortex (Cumming and DeAngelis, 2001; Gonzalez and Perez, 1998; Orban, 2011; Parker, 2007; Welchman, 2016; Welchman and Kourtzi, 2013). We defined activity associated with the derivation of depth or 3D form as those that could be isolated by subtracting brain responses to 2D form images from those to 3D form images (3D-2D contrast). Additionally, we hypothesised that responses in the 3D-2D contrast observed under monocular aperture viewing that were significantly different from other viewing conditions (Vishwanath and Hibbard, 2013) would constitute evidence of activity specifically associated with the impression of monocular stereopsis. We focussed on the analysis of alpha (8-13Hz) and gamma (60-90Hz) frequency ranges within the time window [0-800ms] in both visual and parietal cortex. We were particularly interested in any significant difference during the period before the onset of the fixation colour change [0-800ms] in the high gamma band frequency range of 60 to 90Hz, which has been shown to be related with bottom-up feedforward processes (Bastos et al., 2012; Buschman and Miller, 2007; Engel et al., 2001; Fries, 2015; Wang, 2010).

Data analysis was conducted for only trials with correct responses with a trial number group mean (±standard error [SE]) of 500 ± 3.2 trials remaining (B-3D: 63 ± 0.5; B-2D: 63 ± 0.4; M-3D: 63 ± 0.4; M-2D: 63 ± 0.3; BA-3D: 62 ± 0.5; BA-2D: 62 ± 0.7; MA-3D: 62 ± 0.8; MA-2D: 62 ± 0.7), excluding miss and error trials. Data were subsequently bandpass filtered (EEG: 1-120Hz) using a non-causal Hamming windowed sinc FIR filter (actual cutoff frequency of 0.5 and 120.5Hz at −6dB), re-referenced to average channel values, and then notch filtered (45-55Hz) using a non-causal Hamming windowed sinc FIR filter (actual cutoff frequency of 46 and 54Hz at −6dB) to remove line noise before been epoched into single-trials from −1.5s to 1s relative to the onset of the geometric form image in each trial (EEGLAB, https://sccn.ucsd.edu/eeglab/). The epoched duration of single trials were determined to include sufficient baseline and primary interest of the probe visual cue period. The probe visual cue period (800ms) was believed to be long enough for the expected effects of stereopsis processing to be observable in the EEG data, as based on the previous findings that depth perception is fully processed by around 250-300ms (Spang et al., 2012). In order to achieve cleaner EEG data, around 5-10% rejection rate was aimed at (Delorme et al., 2007). Applying amplitude threshold of −500 to 500uV and applying improbability test with 5SD for single channels, noisy EEG trials and channels that were contaminated with large motion artefacts were detected and removed. This resulted in a group mean (±standard error [SE]) of 442 ± 10.7 trials (86.2% ± 2.1 of the total) remaining (B-3D: 57 ± 1.2; B-2D: 57 ± 1.6; M-3D: 56 ± 1.7; M-2D: 57 ± 1.8; BA-3D: 52 ± 3.2; BA-2D: 53 ± 3.3; MA-3D: 55 ± 1.5; MA-2D: 56 ± 1.2) for further analysis. Independent component analysis of the EEG data (ICA, EEGLAB) was then used to remove eye-blinks/movements (Delorme et al., 2007; Delorme and Makeig, 2004; Jung et al., 2000), with an average of 7 ICs (SE = 1) removed per subject, and data were re-referenced to an average of all non-noisy channels.

Individual, 4-layer (scalp, skull, CSF, & brain) boundary element (BEM) head models were constructed from the MNI standard anatomical image using the Fieldtrip toolbox (http://www.ru.nl/neuroimaging/fieldtrip) (Oostenveld et al., 2011). A Linearly Constrained Minimum Variance (LCMV) beamformer (Robinson and Vrba, 1998; van Drongelen et al., 1996; van Veen et al., 1997) was then employed to separately spatially localize changes in each subject’s alpha (8-13Hz) and gamma (60–90Hz) frequency oscillations (filtered using 2nd order Butterworth filters) between 0s to 0.8s after the geometric image onset. For each subject and frequency band (alpha or gamma), source power during the active (0s to 0.8s) and passive (−0.5s to 0s) time windows, relative to the image onset, were calculated.

Subsequently, pseudo T-statistic (T-statistic) maps were computed as the ratio of the difference in source power between the active and passive windows, divided by the sum of the noise power estimates inherent to the sensors during both active and passive windows (Hillebrand and Barnes, 2005; Robinson and Vrba, 1998).

The minimum peak T-statistic location of the alpha power ERD and maximum peak T-statistic location of the gamma power ERS in the visual and parietal cortex defined the sites of alpha and gamma virtual electrodes (VE). A beamformer estimate of the timecourse of neural activity was then extracted from these two VE locations within each visual and parietal cortex using the entire broadband (1–120 Hz) dataset (for more details, please refer to Brookes et al., 2009, 2008, 2004). Time-frequency spectrograms of alpha and gamma VE data were calculated using a multitaper wavelet approach (Scheeringa et al., 2011). Windows of 0.4s duration were moved across the data in steps of 50ms, resulting in a frequency resolution of 2.5Hz, and the use of seven tapers resulted in a spectral smoothing of ±10Hz. Using the mean of the passive window data as baseline the spectrograms were converted to display change in activity relative to baseline.

Two-sided multiple sample-specific t-tests were used to examine significant differences in gamma ERS and alpha ERD between 3D vs. 2D form images under each viewing condition. Statistical inference was based on a non-parametric cluster-based permutation test (Fieldtrip). This test thresholded the time-frequency t-maps at a value of 1.96 (corresponding to a two-sided t-test with an alpha level of 0.05), resulting in time-frequency clusters. The t-values were summed per cluster and used as the test statistic (Maris, 2012; Maris and Oostenveld, 2007; Nichols and Holmes, 2002). A randomization distribution of this test statistic was determined by randomly exchanging 1000 times, and the p-value was approximated by a Monte Carlo estimate. This was done for all possible permutations given the number of subjects, and for each randomization only the maximal test statistic was retained. An observed cluster was deemed significant if it fell outside the central 95% of this randomization distribution, corresponding to a two-sided random effect test with 5% false-positives, corrected for the multiple comparisons across times and frequencies. The significant spectral-temporal cluster masked the raw time-frequency representation difference. The permutation test has advantages over the Bonferroni correction for multiple comparisons, because the Bonferroni correction assumes that all measures are independent, an assumption that is too strong and weakens the power of the statistical test. In contrast, the permutation test considers the true dependency among all of the measures.

#### Stereopsis rating

Rating data consisted of ratings of strength of stereopsis impression on a self-defined scale from 1 (= binocular viewing) to 5 (= monocular aperture viewing) whilst viewing 3D images. Raw ratings were averaged across four images for each subject. Separate sample t-tests were performed to compare binocular-aperture and monocular against monocular-aperture (reference-point of five) and binocular-aperture and monocular against binocular (reference-point of one). Furthermore, one paired t-test was performed to compare the difference between the binocular-aperture and monocular conditions. In order to control for multiple comparisons, the Bonferroni method was used to adjust the p value for 5 pairwise comparisons.

## Results

### Stereopsis rating

Based on the overall results averaged over all participants, the order in the perceived strength of stereopsis was Binocular (1), Monocular (1.4), Binocular Aperture (4.0) and Monocular aperture (5). The Binocular aperture conditions (4.0 ± 0.2) revealed significantly higher ratings than both Binocular and Monocular conditions (*t*(11) = 18.69, *p* <.05, *t*(11) = 19.33, *p* <.05), but significantly lower ratings compared to monocular aperture viewing, *t*(11) = −6.23, *p* <.05. The monocular condition (1.4 ± 0.1) also had significantly lower ratings compared to the monocular aperture viewing, *t*(11) = −27.59, *p* <.05. The ordering of rating was consistent with those found previously in Vishwanath and Hibbard (2013), though in that study the rating difference between monocular aperture and binocular aperture were larger. There are some significant differences between studies. In the current study, participants did an absolute rating from memory, while in the previous study it was always a relative rating between pairs of conditions, and subjects viewed the reference condition every time before making a rating. Making an absolute judgement on a psychological scale from memory may be harder than making relative judgements.

#### Behavioural data on the visual detection task

Table 1 shows the mean RT and response accuracy data for each viewing and stimulus condition. For the response accuracy, there was a significant main effect of Aperture *F*(1, 11) = 6.77, *p* <. 05, *ηp^2^* = .38, with a higher proportion of correct responses in the No-Aperture condition (98.3% ± 0.4) than the Aperture condition (96.9% ± 0.9). No other effects reached significant levels, all *p*s > .05.

**Table 1.** Mean reaction times (RT) in ms and response accuracy (% Correct). SEs are presented in parentheses.

|  | RT |  | Accuracy |  |
| --- | --- | --- | --- | --- |
|  | No Aperture | Aperture | No Aperture | Aperture |
| Monocular 3D | 461.2 (16.8) | 524.0 (19.5) | 98.8 (0.5) | 96.4 (1.2) |
| Monocular 2D | 471.9 (15.2) | 538.0 (17.4) | 98.4 (0.5) | 96.6 (1.1) |
| Binocular 3D | 441.4 (12.8) | 489.9 (22.9) | 98.2 (0.8) | 97.5 (0.7) |
| Binocular 2D | 444.1 (13.3) | 483.4 (23.0) | 97.9 (0.6) | 97.0 (1.1) |

For reaction time, there was a significant main effect of Type of vision, *F*(1, 11) = 7.61, *p* <.05, *ηp^2^* = .41, with responses to monocular viewing being slower (498.8ms ± 15.3) than to binocular viewing (464.7ms ± 16.0). There was also a main effect of Aperture, *F*(1, 11) = 11.48, *p* <.05, *ηp^2^* = .51, with responses to aperture viewing being slower (508.8ms ± 18.8) than non-aperture viewing (454.7ms ± 13.7). There was a significant interaction between Type of vision and Image dimension, *F*(1, 11) = 13.25, *p* <.05, *ηp^2^* = .55. No other effects reached significant levels, all *p*s > .05.

#### EEG data

##### Alpha Virtual Electrodes (alpha VE)

Figure 3 shows the group average T-statistic map of changes in EEG alpha power during the active window (0-800ms after stimulus onset) compared to the passive window (−500 −0ms pre-stimulus fixation) for the 3D form condition for the visual cortex (left panels) and the parietal cortex (right panels). Corresponding T-statistic maps for the 2D form condition are shown in Supplementary figures S2. Decreases in alpha power (ERD, negative T values) were observed in large areas of the visual cortex under all conditions. After calculating the minimum power locations for the contrast 3D-2D form image T values as alpha VE locations, the mean group alpha VE locations in the visual cortex was found: at [−20, −95, 5] mm [MNI:x,y,z] for monocular, at [5, −95, 5] for binocular, at [−20, −75, 30] for monocular aperture, and at [5, −95, 5] for binocular aperture viewing (see Fig.3, crosshairs, left panels). Similarly, the mean group alpha VE locations in the parietal cortex was found: at [−20, −75, 45] mm [MNI:x,y,z] for monocular, at [−30, −45, 60] for binocular, at [−25, −50, 60] for monocular aperture, and at [−20, −50, 60] for binocular aperture viewing (see Fig.3, crosshairs, right panel).

**Figure 3.**
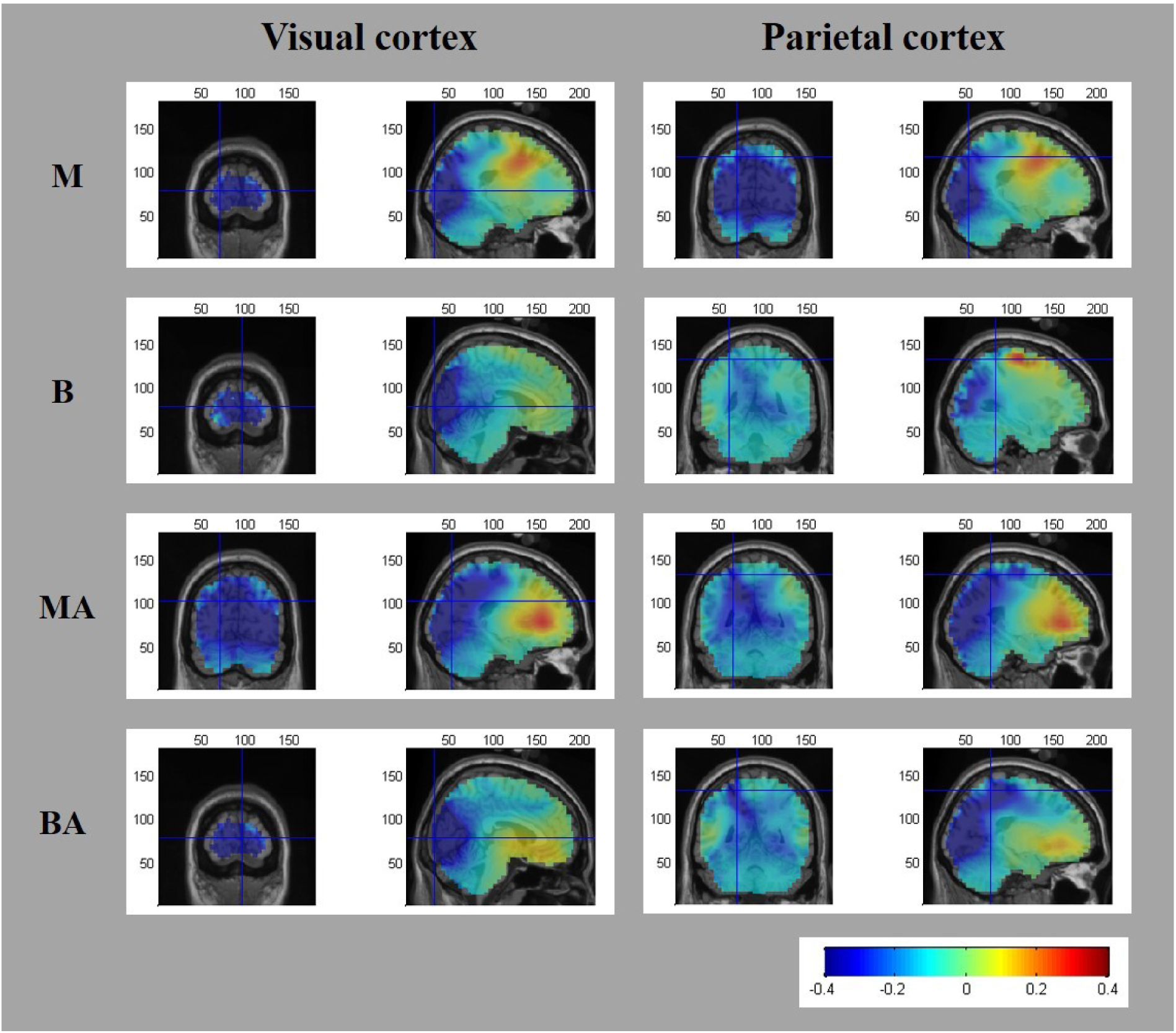
Group average (N=12) T-statistic beamformer maps showing regions exhibiting power decreases in alpha frequency (8-13Hz) during the active window (0-0.8s) as compared with the passive window (−0.5−0s) while viewing 3D form images under 4 different viewing conditions (M: Monocular; B: Binocular; MA: Monocular Aperture; BA: Binocular Aperture). Beamformer maps for 2D form images are in supplementary figures S2. The crosshairs represent the group average alpha VE locations within the visual cortex (left panels) and parietal cortex (right panels) for each viewing condition as determined by the minimum peak power location of 3D-2D T-statistic beamformer maps.

Figure 4 and 5 shows the group mean time-frequency spectrograms measured from alpha VE locations in the visual cortex and parietal cortex respectively for both the active (0 to 0.8s) and passive (−0.5 to 0s) time windows for each viewing condition. During the active window, ERS of gamma band power (60-90Hz) and ERD of alpha (8-13Hz)/ beta (15-30Hz) band power is evident in each viewing condition. Comparison of these TFRs results for each condition with those from Hoogenboom et al. (2010, 2006) and Scheeringa et al. (2016, 2011) shows very similar overall pattern of gamma and alpha/beta responses within the visual cortex measured during visual stimulation.

**Figure 4.**
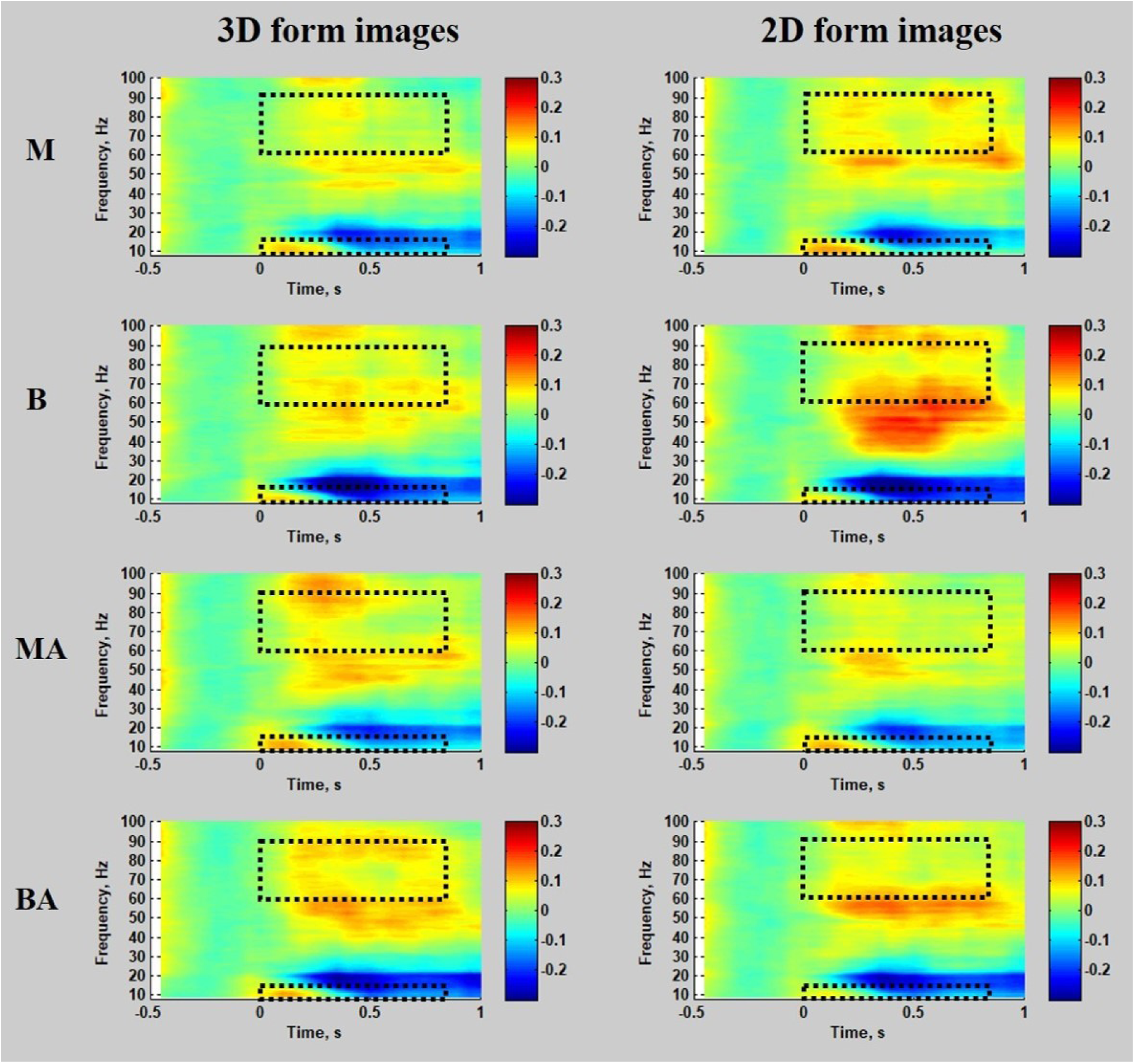
Group mean (N=12) time-frequency spectrograms demonstrating changes in the EEG signal power in alpha ERD VE locations within the visual cortex relative to the passive window (−0.5 to 0s). Time is displayed relative to the initial stimulus onset. Open dashed rectangles represent the a-priori time (0-0.8s) and frequency (alpha: 8-13Hz & high gamma: 60-90Hz) of interest for our analysis. Spectrograms were calculated with frequency resolution of 2.5Hz with spectral smoothing of ±10Hz. Colour bars denote the relative change in power from the average power during the passive window period (baseline measure) of the passive window for each frequency.

**Figure 5.**
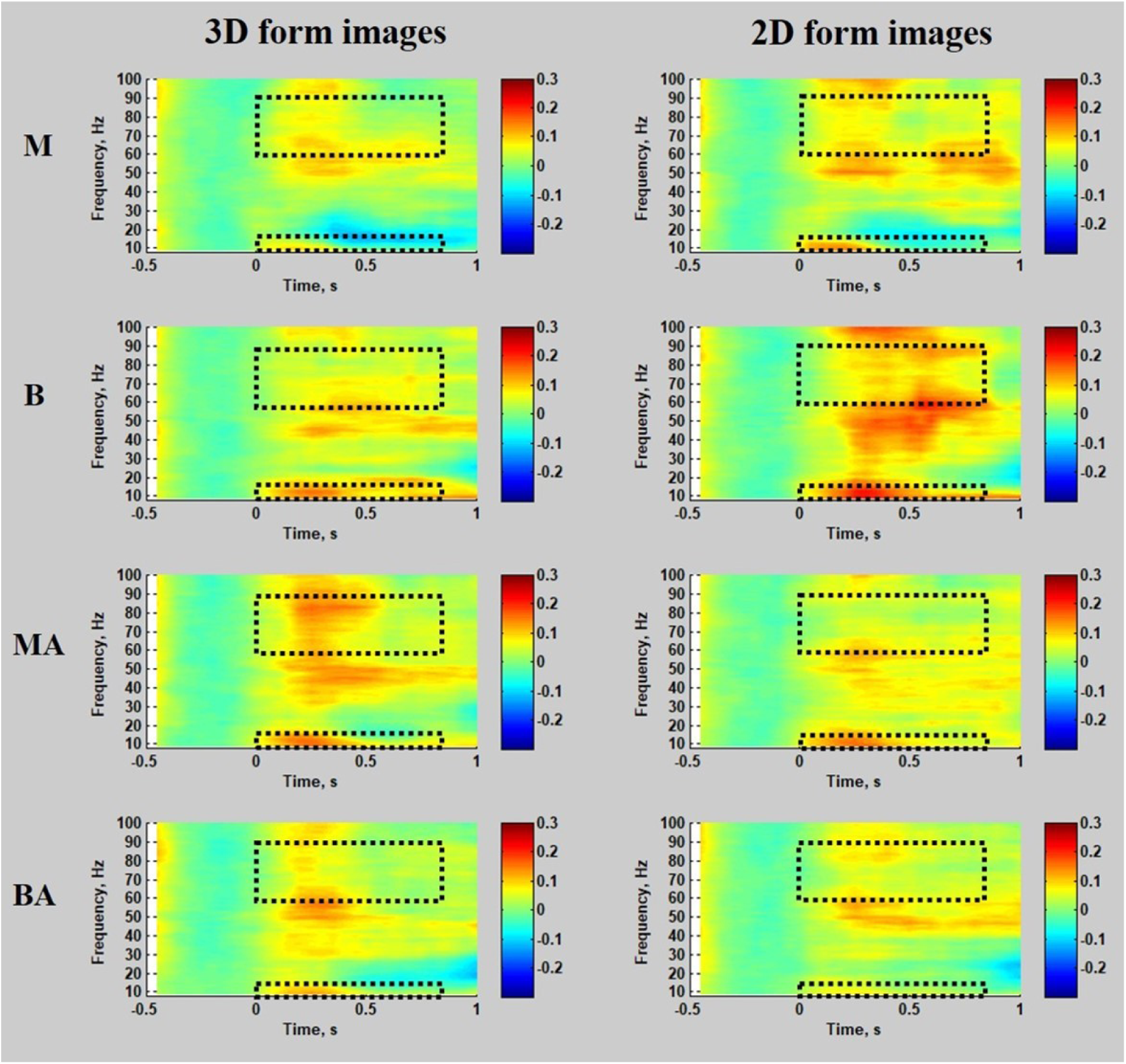
Group mean (N=12) time-frequency spectrograms demonstrating changes in the EEG signal power in alpha ERD VE locations within the parietal cortex relative to the passive window (−0.5 to 0s). Time is displayed relative to the initial stimulus onset. Spectrograms were calculated with frequency resolution of 2.5Hz with spectral smoothing of ±10Hz. Colour bars denote the relative change in power from the average power during the passive window period (baseline measure) of the passive window for each frequency. Open dashed rectangles represent our analysis a-priori interest of time (0-0.8s) and frequency (alpha: 8-13Hz & high gamma: 60-90Hz).

Figure 6 shows the difference between the time-frequency representations (TFRs) of alpha VEs in the visual cortex for 3D and 2D form images (3D-2D) under each viewing condition. After selecting the a-priori time (0-0.8s) and frequency bands (alpha: 8-13Hz & high gamma: 60-90Hz) of interest, cluster-based permutation tests revealed no significant difference between 3D and 2D form images for any of the viewing conditions for the alpha VE locations in the visual cortex (*p* >.05). Figure 7 plots the difference between the TFRs of alpha VEs in the parietal cortex for 3D and 2D form images (3D-2D) under each viewing condition. Fig.7a shows the raw simple differences between the two TFRs. After selecting the a-priori time (0-0.8s) and interest of frequency (alpha: 8-13Hz & high gamma: 60-90Hz) in our data, the cluster-based permutation tests revealed a significant difference in gamma ERS for the contrast 3D - 2D only under the monocular aperture viewing in the alpha VE in the parietal cortex (*p* = .025). Fig.7b shows the significant spectral-temporal cluster of the significant sample-specific t-values (at the corrected alpha-level 0.05, two-sided) within a-priori interest of time (0-0.8s) and alpha frequency (8-13Hz). Fig.7c shows the significant spectral-temporal cluster of the significant sample-specific t-values (at the corrected alpha-level 0.05, two-sided) within a-priori interest of time (0-0.8s) and high gamma frequency (60-90Hz). This indicates that during the interval from 0 to 0.8s, brain responses to 3D form images compared to 2D form images exhibit significantly stronger power in the gamma band (60-90Hz) in the parietal cortex VE locations only in the monocular aperture condition.

**Figure 6.**
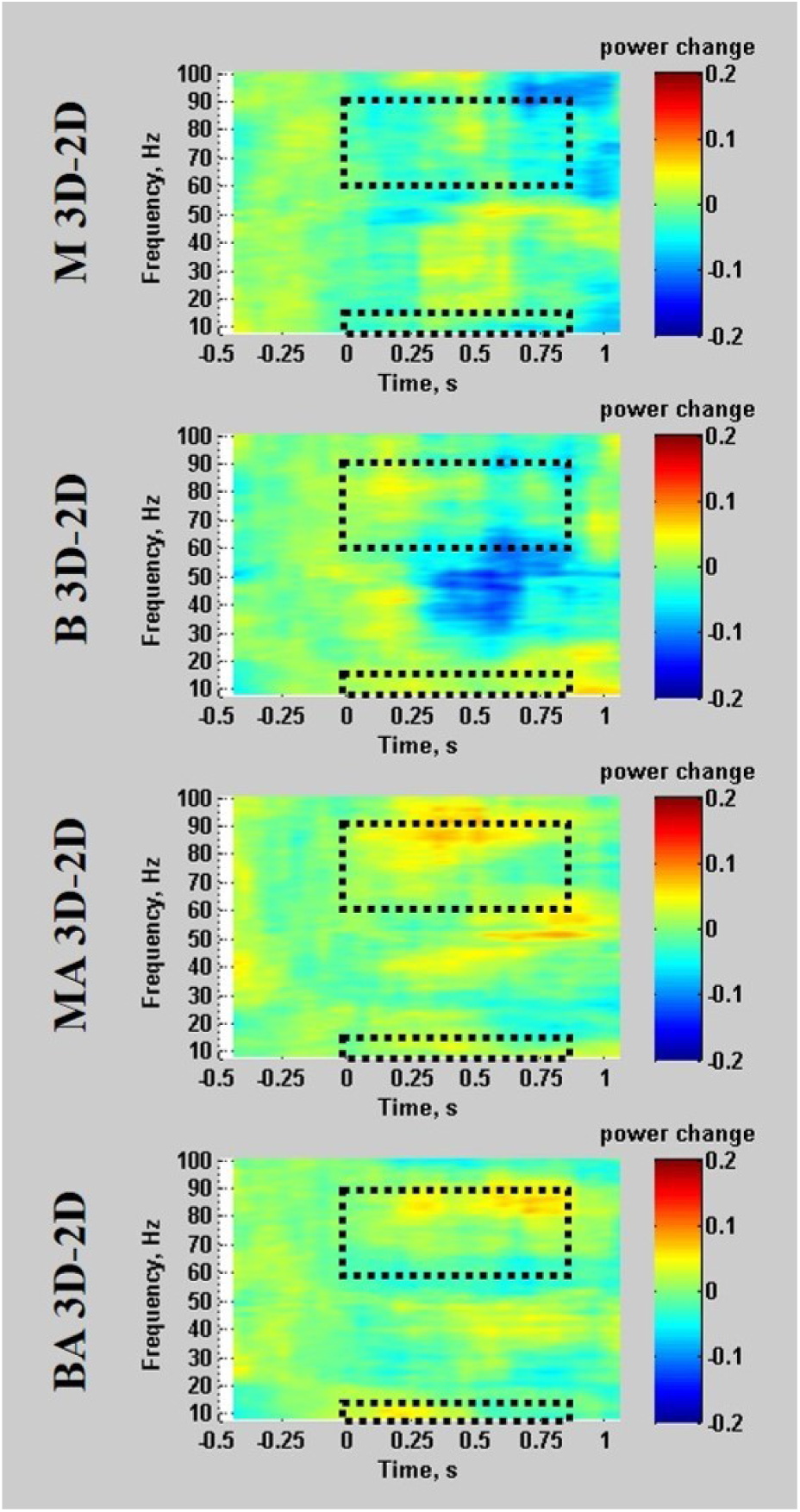
The difference between the time-frequency representations (TFRs) for 3D and 2D form images for alpha VEs in the visual cortex. Simple difference between the two TFRs. Open dashed rectangles represent the a-priori time (0-0.8s) and frequency (alpha: 8-13Hz & high gamma: 60-90Hz) region of interest for our analysis.

**Figure 7.**
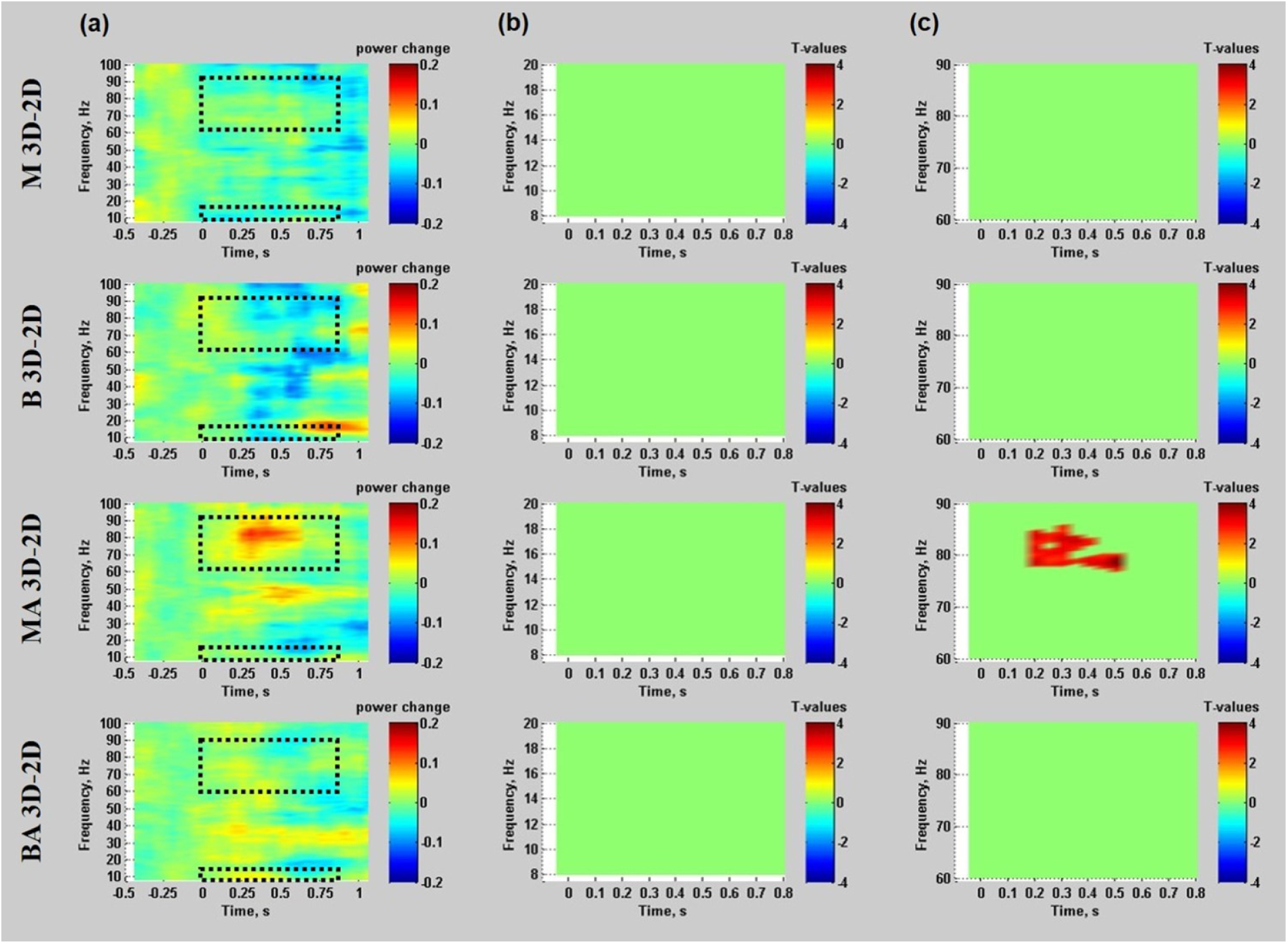
The difference between the time-frequency representations (TFRs) for 3D and 2D form images for alpha VEs in the parietal cortex. (a) Simple difference between the two TFRs. Open dashed rectangles represent the a-priori time (0-0.8s) and frequency (alpha: 8-13Hz & high gamma: 60-90Hz) region of interest for our analysis. (b) Significant spectral-temporal cluster of the significant sample-specific t-values (at the corrected alpha-level 0.05, two-sided) within a-priori interest of time (0-0.8s) and alpha frequency (8-13Hz). (c) Significant spectral-temporal cluster of the significant sample-specific t-values (at the corrected alpha-level 0.05, two-sided) within a-priori interest of time (0-0.8s) and high gamma frequency (60-90Hz).

##### Gamma virtual electrodes (gamma VE)

The group average T-statistic map of changes in EEG gamma power during the active window compared to the passive window are shown in supplementary Figure S3 and S4. Increases in gamma power (ERS, positive T values) were observed in the visual cortex under all conditions. After calculating the maximum power locations of 3D-2D form image T values as gamma VE locations, the mean group gamma VE locations in the visual cortex was found: at [0, −90, 0] mm [MNI:x,y,z] for monocular, at [−30, −85, 0] for binocular, at [−45, −70, 5] for monocular aperture, and at [5, −55, 10] for binocular aperture viewing (see Fig S3 crosshairs). The mean group gamma VE locations in the parietal cortex was found: at [25, −60, 65] mm [MNI:x,y,z] for monocular, at [−50, −20, 35] for binocular, at [−45, −65, 40] for monocular aperture, and at [−55, −50, 45] for binocular aperture viewing (see Fig.S4, crosshairs). However, no significant differences in alpha or gamma band power for the 3D - 2D viewing contrast at either the visual or parietal gamma VE locations was found for the a-priori time (0-0.8s) and frequency (alpha: 8-13Hz & high gamma: 60-90Hz) of interest using the cluster-based permutation tests (all *p* > .05). Relevant plots of time-frequency representations (TFRs) of gamma VEs in the visual and parietal cortex for each viewing condition as well as differences between 3D and 2D TFRs are shown in Supplementary figures (S5, S6 and S7).

## Discussion

In this study, we examined changes in broad neural activity in occipital and parietal cortex when two types of stimulus (2D vs. 3D form) were viewed under four different viewing conditions. Our aim was to determine if we could identify a dissociable neural signature in the specific stimulus-viewing condition (monocular aperture viewing) that generates the qualitative impression of stereopsis in the absence of binocular disparity (Ames, 1925; Koenderink, 1998; Michotte, 1991, 1948; Schlosberg, 1941; Vishwanath and Hibbard, 2013; da Vinci, cited in Wade et al., 2001; Wheatstone, 1838). Specifically we measured EEG activity when subjects viewed 3D and 2D form images under four different viewing conditions (Monocular, Binocular, Monocular Aperture, Binocular Aperture). The results of the behavioural rating task confirmed that the most compelling qualitative impression of stereopsis was experienced for the 3D form images under the monocular aperture viewing condition, consistent with previous results (Vishwanath and Hibbard, 2013). From our analysis of EEG oscillatory activity at visual and parietal cortex virtual electrode (VE) locations derived from both alpha and gamma band activity during the epoch of a priori interest (0-800ms after image onset), the only statistically significant difference we found in oscillatory activity subtracting activation for 2D form images from 3D form images was in gamma (60-90Hz) event-related synchronization (ERS) at the alpha virtual electrode location in parietal cortex under the monocular aperture viewing. There were no statistically significant differences in EEG gamma or alpha band activity between 3D and 2D conditions for visual cortex VE locations in any of the viewing conditions. We also did not find any significant difference in gamma band activity for the parietal cortex gamma band VE location. Taken together, our results provide initial evidence suggestive of dissociable neural activity specifically linked to processes that underlie the phenomenological visual experience associated with stereopsis, as distinguished from disparity processing per se.

## Alpha vs gamma VE

The significant difference in gamma synchronization between 3D and 2D images that we observed under monocular aperture viewing was obtained at the alpha band virtual electrode (VE) location. Alpha and gamma band activities are thought to reflect different but related processes (Buschman and Miller, 2007; Buzsaki and Draguhn, 2004; Colgin et al., 2009; Fries, 2009; Jensen et al., 2014; Klimesch, 1999; Pfurtscheller et al., 1996; Pfurtscheller and Lopes da Silva, 1999; Singer and Gray, 1995), and the alpha VE and gamma VE locations are typically co-located (Ball et al., 2008; Cheyne et al., 2008; Cheyne and Ferrari, 2013; Cheyne, 2013; Donner and Siegel, 2011; Hoogenboom et al., 2006; Muthukumaraswamy, 2013, 2010; Muthukumaraswamy and Singh, 2013; Scheeringa et al., 2011; Uji et al., 2018). One question regarding our results might be why we found no significant effects of gamma ERS at the parietal gamma VE location itself.

Gamma source localization is more susceptible to reduced signal-to-noise ratios (SNRs) due to suboptimal data processing than alpha source localization. While the alpha power change is sustained over longer periods of time and typically observable in EEG with robust SNR, gamma power change is more short lived and has low SNR in the EEG signal (Darvas et al., 2010; Miller et al., 2007). Our study design precluded recording a large number of trials within a workable experimental session duration, and thus we may not have achieved the same level of SNRs required for gamma band source localization that has been achieved in previous studies which typically record in excess of 100 trials per condition (Ball et al., 2008; Cheyne et al., 2008; Cheyne and Ferrari, 2013; Cheyne, 2013; Hoogenboom et al., 2006; Muthukumaraswamy, 2013, 2010; Muthukumaraswamy and Singh, 2013; Scheeringa et al., 2011; Uji et al., 2018); more trials leads to better SNRs (e.g., 120 trials using individual BEM head model, see details Uji et al., 2018). Furthermore, in this study, we used the MNI standard anatomical image and standard EEG electrode locations to construct the BEM head model estimating the neural activities in the brain instead of using individual anatomical brain images and digitizing the EEG electrodes locations (see Uji et al., 2018). The latter approach can improve source localization reliability and SNR (Brookes et al., 2008; van Drongelen et al., 1996; van Veen et al., 1997; Wan et al., 2008). These limiting factors likely negatively impacted on the sensitivity and specificity with which we were able to identifying the optimal gamma VE location. We are therefore more confident of the derived alpha VE locations particularly because previous studies that have had better SNRs (number of trials) and/or use of individual anatomical brain images indicate that alpha and gamma VE locations are spatially co-located (Ball et al., 2008; Cheyne et al., 2008; Cheyne and Ferrari, 2013; Cheyne, 2013; Donner and Siegel, 2011; Hoogenboom et al., 2006; Muthukumaraswamy, 2013, 2010; Muthukumaraswamy and Singh, 2013; Scheeringa et al., 2011; Uji et al., 2018).

## Location of VEs in comparison to previous studies

The alpha virtual electrode location where we found the significant difference in activity was at [−25, −50, 60] mm [MNI:x,y,z] in the parietal cortex. Previous neuroimaging studies on depth from binocular disparity under stimulus conditions generating binocular stereopsis have implicated the parietal cortex (Chandrasekaran et al., 2007; Durand et al., 2009; Georgieva et al., 2009; Minini et al., 2010; Tsao et al., 2003). Specifically, Durand et al. (2009) demonstrated that human anterior intraparietal sulcus (IPS) (Dorsal IPS medial at [−22, −62, 56] mm [MNI:x,y,z]; dorsal IPS anterior at [−30, −50, 64] mm) was activated when processing the 3D structure, whereas the posterior IPS at [−24, −82, 32] mm was activated when processing location in 3D space. Although EEG has lower spatial resolution than fMRI (Bandettini, 2009; Buzsáki et al., 2012; Cohen, 2017; Sejnowski et al., 2014), it is interesting that the alpha virtual electrode (VE) location in the parietal cortex we found was located at [−25, −50, 60] mm in the posterior parietal cortex close to anterior IPS. This is also consistent with the recent finding of differential anterior IPS activation during the viewing phase of visually guided grasping before the movement execution comparing real vs. pictured objects (Freud et al., 2018). They suggested that the “realness” of the target object differentiated the anterior IPS activation. Although they did not detail what visual attributes, cues or visual processing they take to constitute realism, the findings do provide support for our interpretation. Our working assumption, based on data on perceptual phenomenology of 3D perception, is that the key phenomenological characteristics of object solidity/tangibility and the sense of immersive negative space that are characteristic of stereopsis is what primarily underlies the overall visual sensation of realness associated with real and stereoscopic scenes (Vishwanath, 2014).

## Parietal vs. extrastriate regions in generating stereopsis

Accumulating evidence from neuroimaging studies on binocular disparity have demonstrated that the 3D information is processed in the dorsal stream extending beyond the primary visual cortex to the parietal cortex (Chandrasekaran et al., 2007; Durand et al., 2009; Georgieva et al., 2009; Minini et al., 2010; Tsao et al., 2003). In our study, the monocular aperture viewing of pictorial images did not need to integrate disparity or motion defined depth signals, such that the relative activation of dorsal extrastriate regions implicated in disparity or motion processing (i.e. V3A, V3B/KO, MT and V7) might not have shown heightened activity. Our data do not necessarily imply that dorsal extrastriate regions in the visual cortex (i.e. V3A, V3B/KO, MT and V7) are not involved in processing of representation that underlie the qualitative impression of stereopsis. EEG source localizations still have significant limitations in spatial resolution as compared to fMRI (Brookes et al., 2008; Gross, 2016; Wan et al., 2008), and precise and optimal EEG source localization is likely to be difficult in the visual cortex, which has more complicated structure and layers, and required use of retinotopic mapping in fMRI (Backus et al., 2001; Bridge and Parker, 2007; Cottereau et al., 2011; Durand et al., 2009; Goncalves et al., 2015; Minini et al., 2010; Preston et al., 2008). However, despite the limitations of the fairly board approach we employed to detect a dissociable neural signature of monocular stereopsis, our data of significant gamma ERS within the parietal cortex for the monocular aperture condition, taken together with recent evidence for processes in anterior IPS in the parietal cortex for real vs. pictured objects (Freud et al., 2018), provide converging evidence that the visual impressions associated with stereopsis are likely associated with visual encoding in later stages of processing in the dorsal stream.

## Potential Confounds of viewing and stimulus conditions

We focussed on contrasts between 3D and 2D form images for the four viewing conditions in order to eliminate several possible confounding factors that could have interacted with the presence of the qualitative impression of stereopsis. For example, our result cannot be attributed to either lack of binocularity or the presence of the aperture in the monocular aperture condition (MA) because (1) other conditions in which there was no disparity (monocular viewing without aperture) or where there was an aperture (Binocular Aperture viewing) did not generate the same effect. Moreover, the results for each viewing condition was obtained by subtracting activity between stimulus conditions (3D and 2D) for which all aspects other than the depicted 3D structure (such as disparity content, field of view, presence of aperture) was the same. This argues against the interpretation that the significant difference in gamma power in the 3D-2D contrast for the MA viewing condition was due to a generic idiosyncratic variable (e.g., field of view, disparity content) of monocular-aperture viewing independent of image content. Any such generic effect would be present for both the 3D and 2D stimulus conditions and would effectively be subtracted out in the contrast. Moreover the effect cannot be ascribed to generic differences in the image content (e.g., orientation content, differences in local luminance distributions) of the two stimulus types (3D, 2D) since any low-level differences between these two stimulus types did not yield similar differential effects in gamma activity for viewing conditions other than MA viewing. Also, such low-level effects would most likely reveal differences in neural activity in visual rather than parietal cortex (Cumming and DeAngelis, 2001; Gonzalez and Perez, 1998; Orban, 2011; Parker, 2007; Sakata et al., 2005; Welchman, 2016; Welchman and Kourtzi, 2013).

## Behavioural data, motor preparation and attention

The aperture conditions (monocular and binocular) showed a small but statistically significant decrease in detection accuracy compared to the no-aperture conditions. We also found that reaction times were slightly faster for the binocular condition compared to the monocular conditions overall, and the presence of the aperture caused small increase in reaction time. These could have arisen due to the limited visual field of the aperture and the lack of stimulus additivity in the monocular as opposed to the binocular condition. However, the results we obtained from the stimulus contrast in parietal cortex cannot plausibly be attributed to differences in task performance under monocular aperture viewing, as any behavioural effects specific to detection due to generic visual properties of viewing monocularly through an aperture should have affected the 3D and 2D condition equally, and therefore be subtracted out in the contrast. Although there was a small difference in RT between the 3D and 2D form images under monocular aperture viewing, this difference was not significant [mean RTs (± SE) for MA-3D (524.0ms ± 19.5) and MA-2D (538.0ms ± 17.4)]. Moreover, previous studies examining Monkey V4 (Schoffelen et al., 2005), human visual (Hoogenboom et al., 2010) and motor cortex (Womelsdorf et al., 2006) consistently show that stronger gamma-band activity predicted shorter behavioural response times, which is opposite to the potential interpretation that the increase in gamma power that we found in the parietal VE was due to the insignificanty longer behavioural response times in the MA-3D condition.

The higher gamma ERS observed in the parietal cortex VE location under the monocular aperture viewing could not be plausibly related with preparation for the movement execution as such preparation should have been common to all conditions and would have been subtracted out in the contrasts (3D-2D). Like us, Freud et al. (2018), who also found greater activation in anterior IPS during the planning phase of visually guided grasping of real vs. pictured objects before the movement execution, attribute the activity to the visual perception during response preparation, rather than the response preparation itself.

Could the results be attributed to differential attentional state either due to generic differences in viewing condition or stimulus content? This is unlikely. First, we controlled allocation of attention across all stimulus and viewing condition by utilizing a detection task that required constant monitoring of a small visual point for the full duration of the time of EEG data analysis. Second, any differences arising from viewing conditions (e.g. presence of the aperture) should be the same for the 3D and 2D form condition and therefore in effect subtracted out in our contrasts.

## Conclusion

This study is the first to our knowledge to examine whether there is dissociable neural activity linked to the qualitative impression of stereopsis (solidity, tangibility, negative space and realness) that is not confounded with activity underlying disparity processing or the perception of pictorial 3D form and depth relations. We utilised a fairly broad approach by analysing gross neural activity as measured via EEG oscillatory responses in the alpha and gamma domain. Our results provide the first glimpse that such distinct activity may indeed exist and that it is likely localized to parietal regions processing visuo-motor transformations. This would be consistent with claims that the visual impression of stereopsis is associated with the conscious awareness of the capacity to manipulate 3D objects (Michotte, 1948). Our results are consistent with other recent work in fMRI that has also implicated regions of the parietal cortex in differentiating visual perception guiding movements to either real or pictured objects (Freud et al., 2018). Future research will need to provide further confirmatory evidence and determine if the representation and processes that bring about the experience of stereopsis and realness are the same regardless of the source of the depth signal (binocular disparity, motion parallax or pictorial cues) and also identify the specific stage of transformation of visual information that underlies this central phenomenological aspect of human 3D space perception.

## Acknowledgements

We would like to thank Justin Ales for helpful comments on earlier versions of the manuscript.

## Supplementary Materials

**Figure S1.**
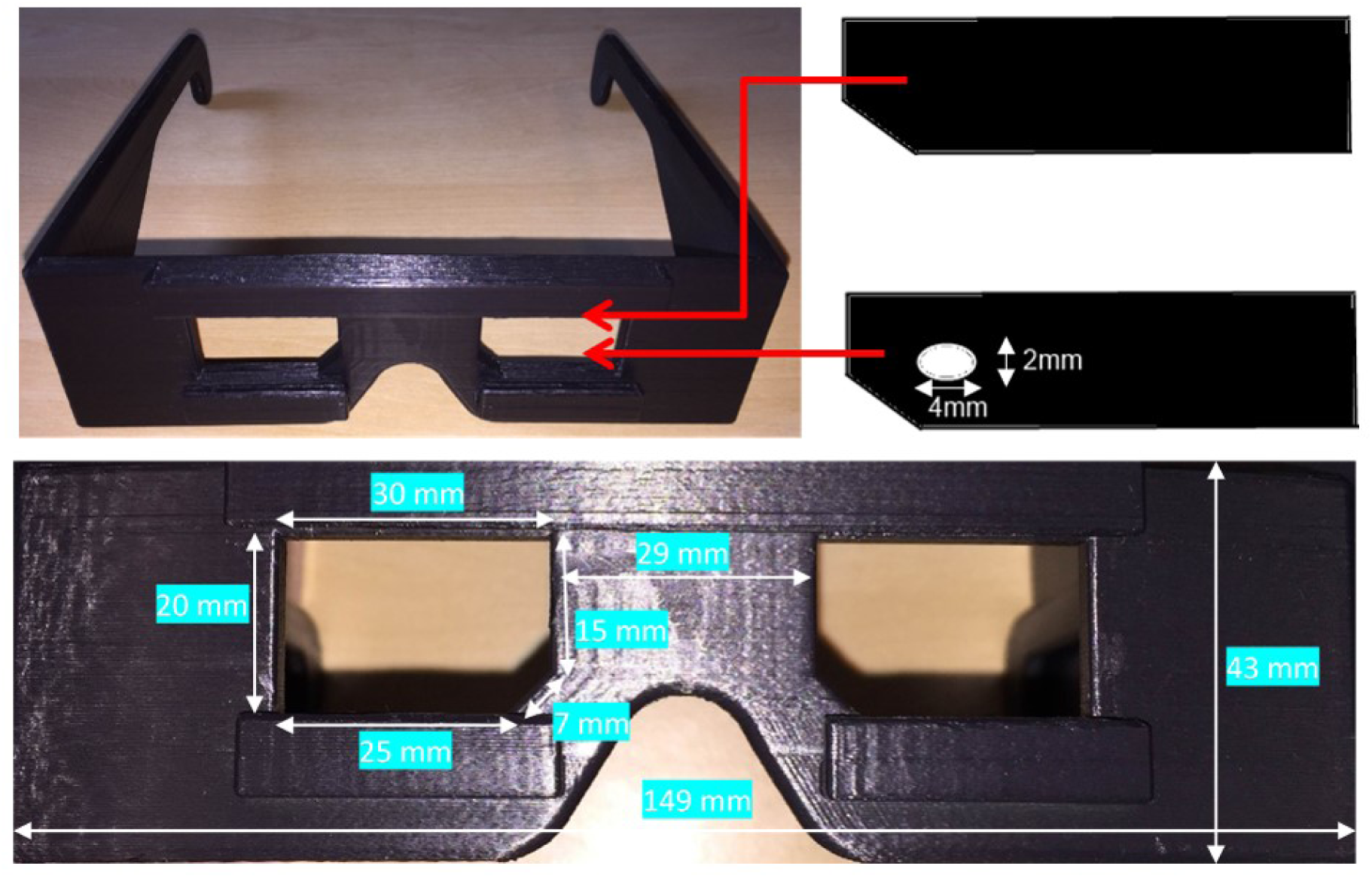
Plastic spectacles worn by participants during the experiment. The spectacles were placed over the EEG cap and over spectacles for corrected-to-normal vision. The temples tapered in thickness, reducing the peripheral visual field. Cardboard slides were inserted to create monocular and aperture viewing-conditions.

**Figure S2.**
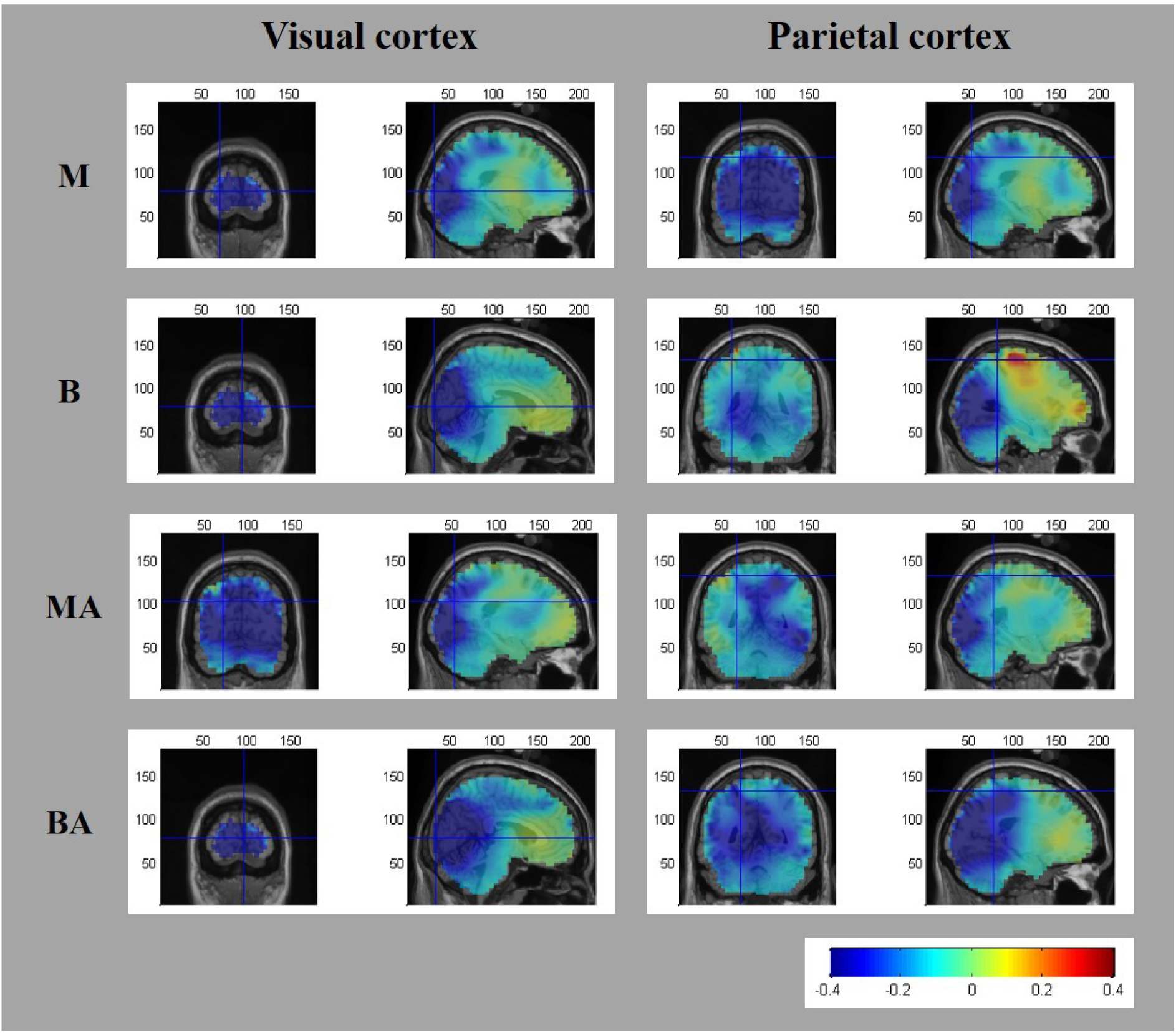
Group average (N=12) T-statistic beamformer maps showing regions exhibiting power decreases in alpha frequency (8-13Hz) during the active window (0-0.8s) as compared with the passive window (−0.5-0s) while viewing 2D form images under 4 different viewing conditions (M: Monocular; B: Binocular; MA: Monocular Aperture; BA: Binocular Aperture). The crosshairs represent the group average alpha VE locations within the visual cortex (left) and parietal cortex (right) for each viewing condition as determined by the minimum peak power location of 3D-2D T-statistic beamformer maps.

**Figure S3.**
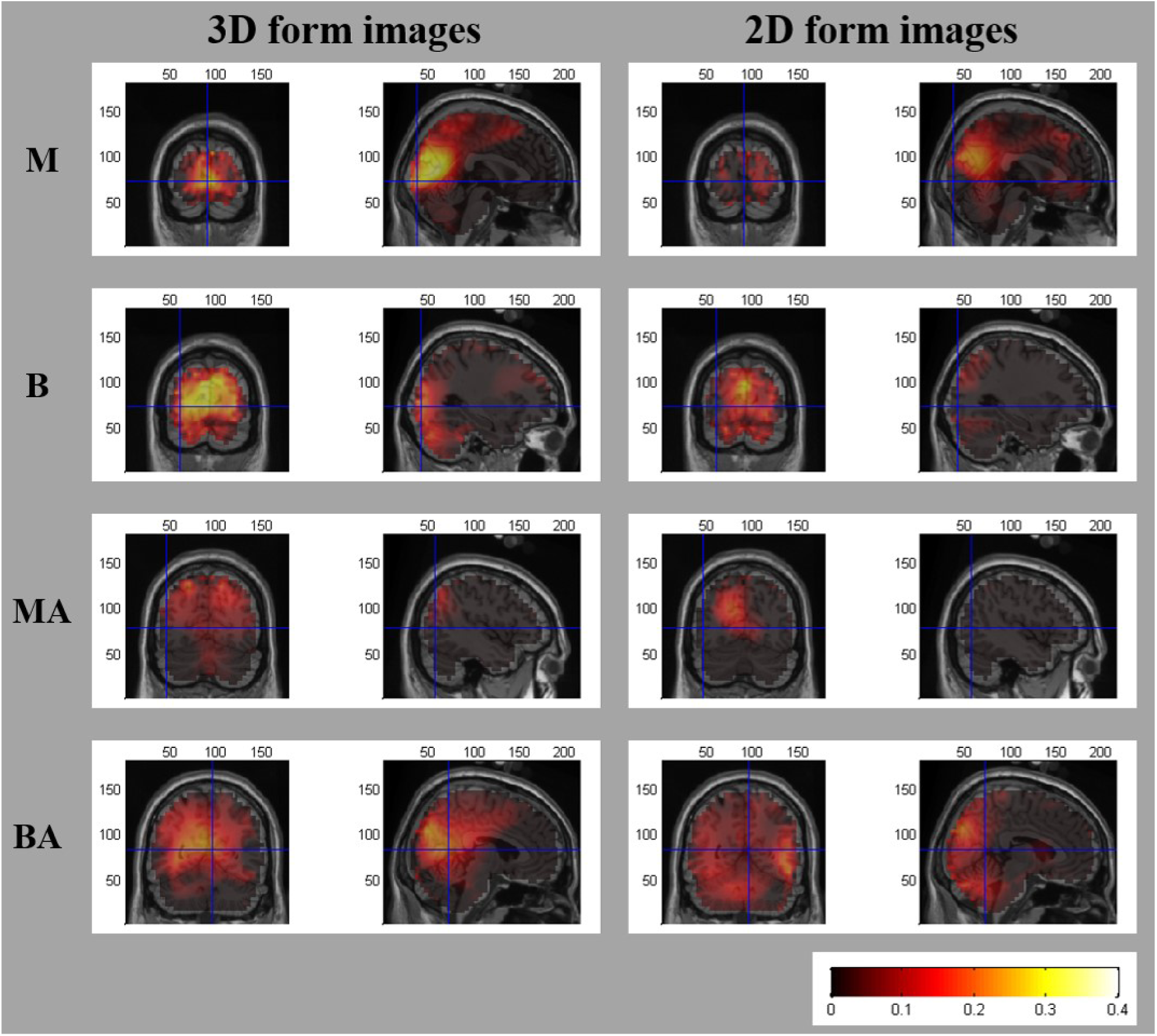
Group average (N=12) T-statistic beamformer maps showing regions exhibiting power increases in gamma frequency (60-90Hz) during the active window (0-0.8s) as compared with the passive window (−0.5-0s) while viewing either 3D form (left column) or 2D form images (right column) under 4 different viewing conditions (M: Monocular; B: Binocular; MA: Monocular Aperture; BA: Binocular Aperture). The crosshairs represent the group average gamma VE locations within the visual cortex for each viewing condition as determined by the maximum peak power location of 3D-2D T-statistic beamformer maps.

**Figure S4.**
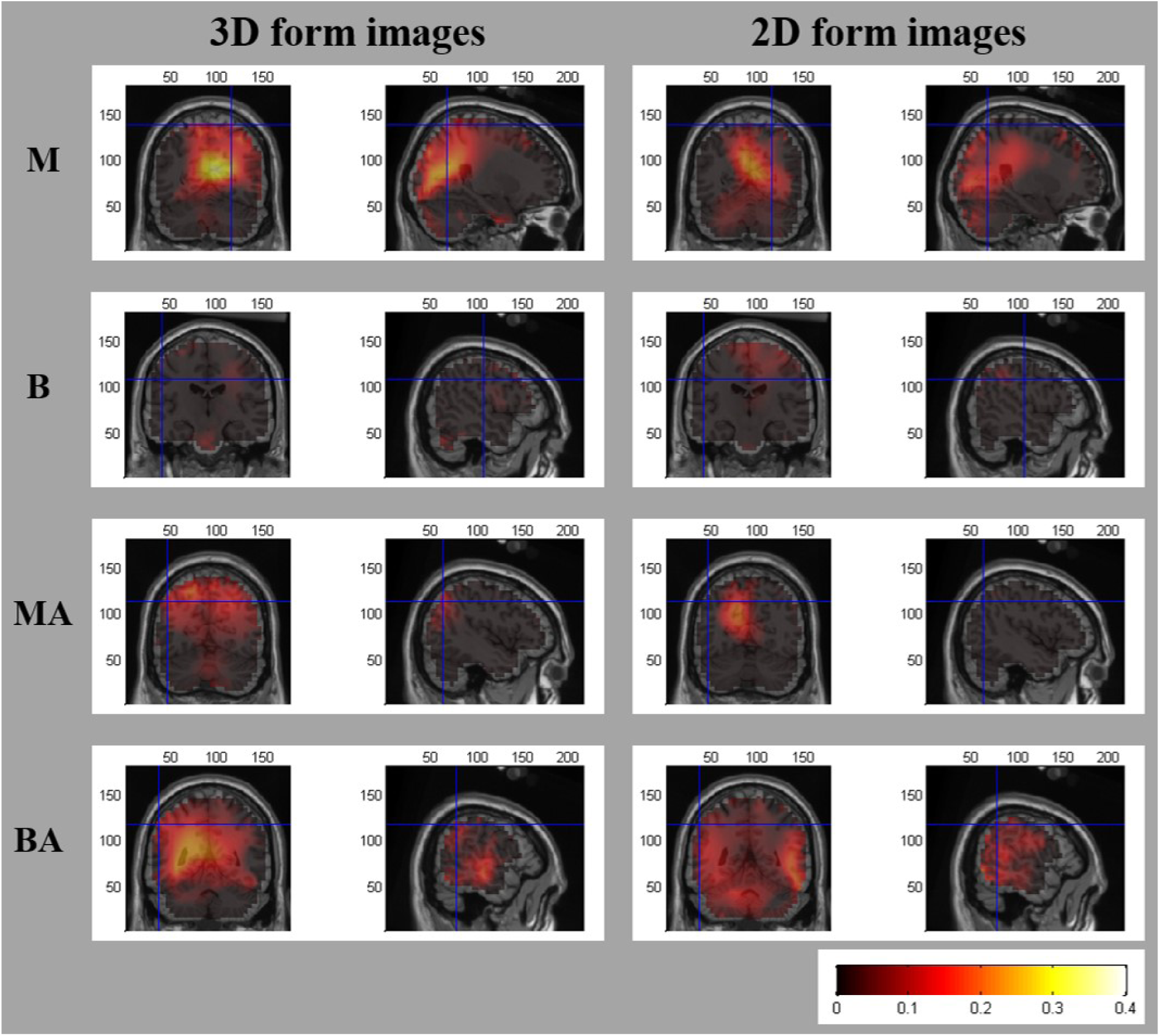
Group average (N=12) T-statistic beamformer maps showing regions exhibiting power increases in gamma frequency (60-90Hz) during the active window (0-0.8s) as compared with the passive window (−0.5-0s) while viewing either 3D form (left column) or 2D form images (right column) under 4 different viewing conditions (M: Monocular; B: Binocular; MA: Monocular Aperture; BA: Binocular Aperture). The crosshairs represent the group average gamma VE locations within the parietal cortex for each viewing condition as determined by the maximum peak power location of 3D-2D T-statistic beamformer maps.

**Figure S5.**
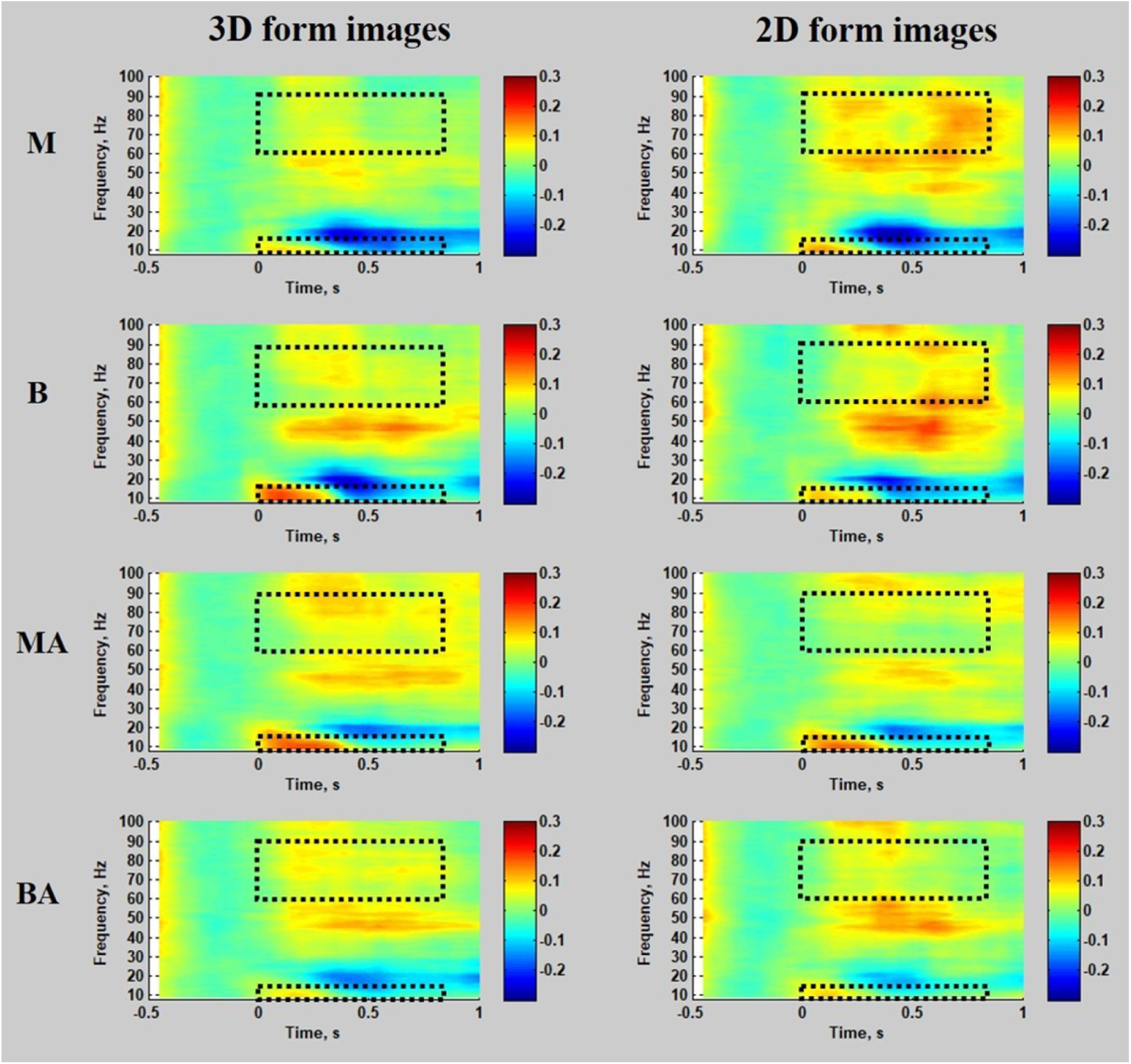
Group mean (N=12) time-frequency spectrograms demonstrating changes in the EEG signal power in gamma ERS VE locations within the visual cortex relative to the passive window (−0.5 to 0s). Time is displayed relative to the initial stimulus onset. Spectrograms were calculated with frequency resolution of 2.5Hz with spectral smoothing of ±10Hz. Colour bars denote the relative change in power from the average power during the passive window period (baseline measure) of the passive window for each frequency. Open dashed rectangles represent our analysis a-priori interest of time (0-0.8s) and frequency (alpha: 8-13Hz & high gamma: 60-90Hz).

**Figure S6.**
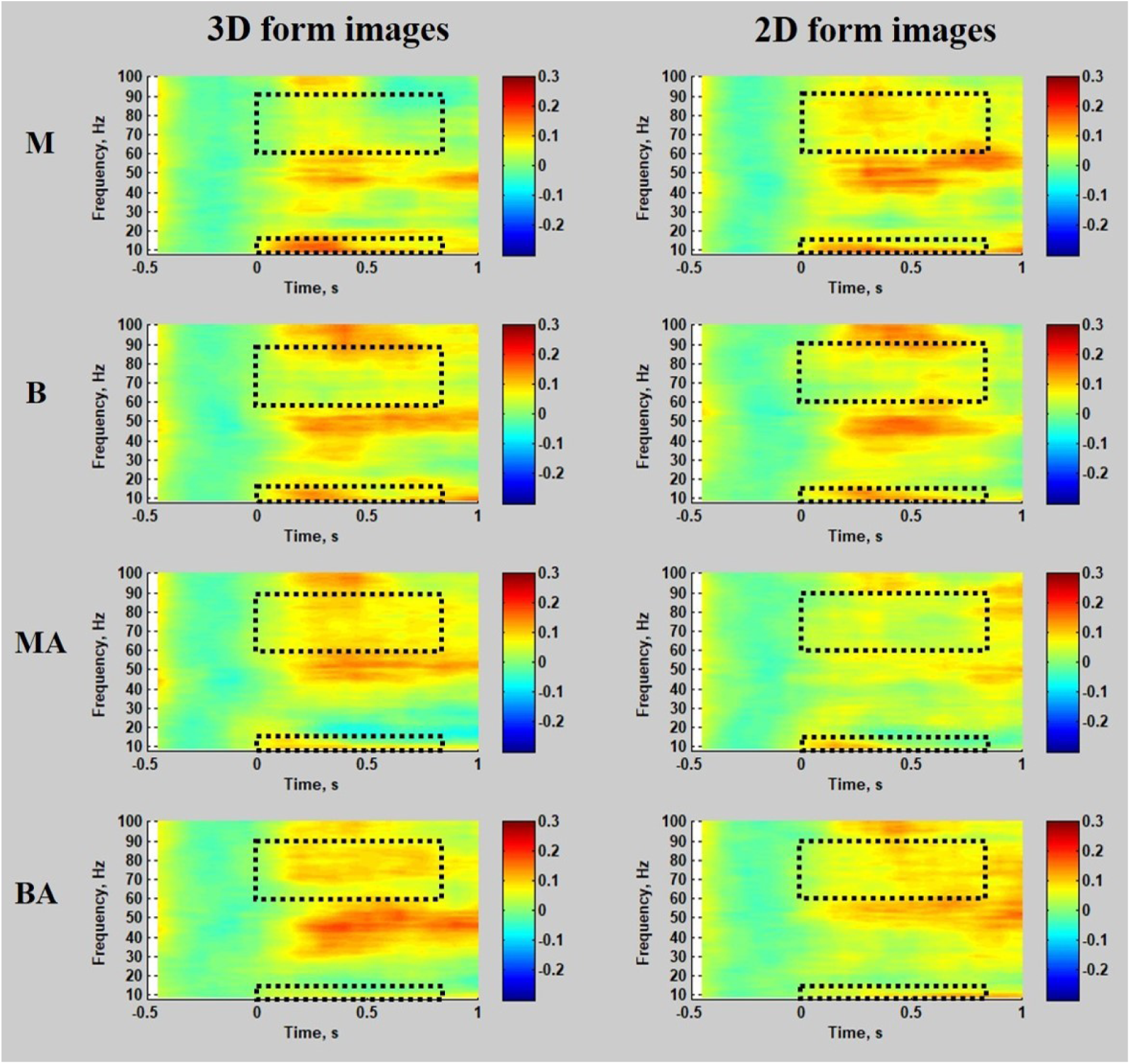
Group mean (N=12) time-frequency spectrograms demonstrating changes in the EEG signal power in gamma ERS VE locations within the parietal cortex relative to the passive window (−0.5 to 0s). Time is displayed relative to the initial stimulus onset. Spectrograms were calculated with frequency resolution of 2.5Hz with spectral smoothing of ±10Hz. Colour bars denote the relative change in power from the average power during the passive window period (baseline measure) of the passive window for each frequency. Open dashed rectangles represent our analysis a-priori interest of time (0-0.8s) and frequency (alpha: 8-13Hz & high gamma: 60-90Hz).

**Figure S7.**
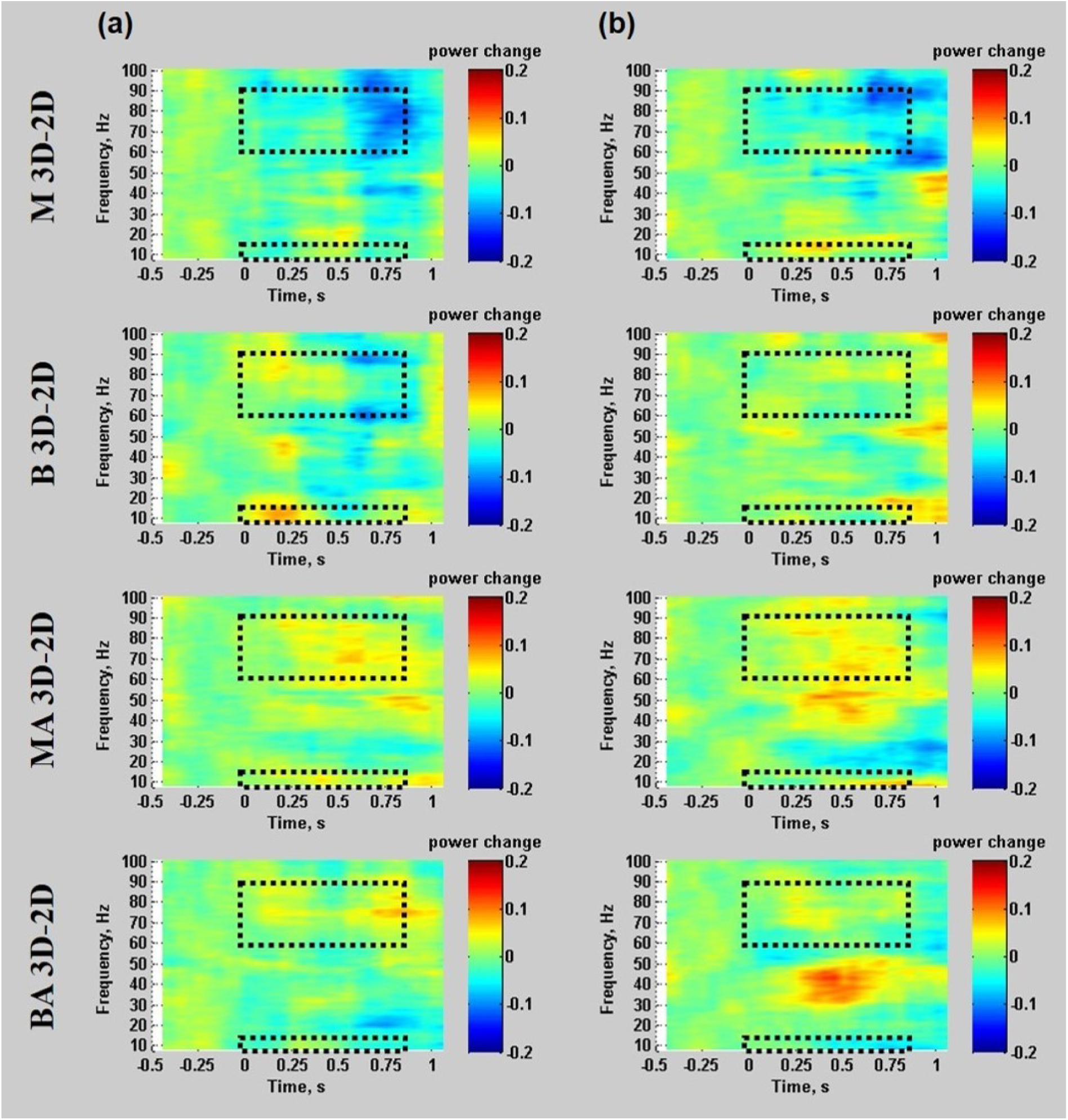
The difference between the time-frequency representations (TFRs) for 3D and 2D form images for gamma VEs in the visual cortex (a) and parietal cortex (b). Open dashed rectangles represent our analysis a-priori interest of time (0-0.8s) and frequency (alpha: 8-13Hz & high gamma: 60-90Hz).

## References

Ames, A., 1925. The Illusion of Depth from Single Pictures. J. Opt. Soc. Am. 10, 137–148. https://doi.org/10.1364/JOSA.10.000137

Anzai, A., DeAngelis, G.C., 2010. Neural computations underlying depth perception. Curr. Opin. Neurobiol. 20, 367–375. https://doi.org/10.1016/j.conb.2010.04.006

Backus, B.T., Fleet, D.J., Parker, A.J., Heeger, D.J., 2001. Human Cortical Activity Correlates With Stereoscopic Depth Perception. J. Neurophysiol. 86, 2054–2068. https://doi.org/10.1152/jn.2001.86.4.2054

Ball, T., Demandt, E., Mutschler, I., Neitzel, E., Mehring, C., Vogt, K., Aertsen, A., Schulze-Bonhage, A., 2008. Movement related activity in the high gamma range of the human EEG. Neuroimage 41, 302–310. https://doi.org/10.1016/j.neuroimage.2008.02.032

Ban, H., Welchman, A.E., 2015. fMRI Analysis-by-Synthesis Reveals a Dorsal Hierarchy That Extracts Surface Slant. J. Neurosci. 35, 9823–9835. https://doi.org/10.1523/JNEUROSCI.1255-15.2015

Bandettini, P.A., 2009. What’s New in Neuroimaging Methods? Ann. N. Y. Acad. Sci. 1156, 260–293. https://doi.org/10.1111/j.1749-6632.2009.04420.x

Barlow, H.B., Blakemore, C., Pettigrew, J.D., 1967. The neural mechanism of binocular depth discrimination. J. Physiol. 193, 327–342. https://doi.org/10.1113/jphysiol.1967.sp008360

Bastos, A.M., Litvak, V., Moran, R., Bosman, C.A., Fries, P., Friston, K.J., 2015. A DCM study of spectral asymmetries in feedforward and feedback connections between visual areas V1 and V4 in the monkey. Neuroimage 108, 460–475. https://doi.org/10.1016/j.neuroimage.2014.12.081

Bastos, A.M., Usrey, W.M., Adams, R.A., Mangun, G.R., Fries, P., Friston, K.J., 2012. Canonical Microcircuits for Predictive Coding. Neuron 76, 695–711. https://doi.org/10.1016/j.neuron.2012.10.038

Bauer, M., Oostenveld, R., Peeters, M., Fries, P., 2006. Tactile Spatial Attention Enhances Gamma-Band Activity in Somatosensory Cortex and Reduces Low-Frequency Activity in Parieto-Occipital Areas. J. Neurosci. 26, 490–501. https://doi.org/10.1523/JNEUROSCI.5228-04.2006

Bauer, M., Stenner, M.-P., Friston, K.J., Dolan, R.J., 2014. Attentional Modulation of Alpha/Beta and Gamma Oscillations Reflect Functionally Distinct Processes. J. Neurosci. 34, 16117–16125. https://doi.org/10.1523/JNEUROSCI.3474-13.2014

Behrmann, M., Geng, J.J., Shomstein, S., 2004. Parietal cortex and attention. Curr. Opin. Neurobiol. 14, 212–217. https://doi.org/10.1016/j.conb.2004.03.012

Beringer, J., 1992. Timing accuracy of mouse response registration on the IBM microcomputer family. Behav. Res. Methods, Instruments, Comput. 24, 486–490. https://doi.org/10.3758/BF03203585

Binkofski, F., Dohle, C., Posse, S., Stephan, K.M., Hefter, H., Seitz, R.J., Freund, H.J., 1998. Human anterior intraparietal area subserves prehension: A combined lesion and functional MRI activation study. Neurology 50, 1253–1259. https://doi.org/10.1212/WNL.50.5.1253

Bridge, H., Parker, A.J., 2007. Topographical representation of binocular depth in the human visual cortex using fMRI. J. Vis. 7, 15. https://doi.org/10.1167/7.14.15

Brookes, M.J., Gibson, A.M., Hall, S.D., Furlong, P.L., Barnes, G.R., Hillebrand, A., Singh, K.D., Holliday, I.E., Francis, S.T., Morris, P.G., 2004. A general linear model for MEG beamformer imaging. Neuroimage 23, 936–946. https://doi.org/10.1016/j.neuroimage.2004.06.031

Brookes, M.J., Mullinger, K.J., Stevenson, C.M., Morris, P.G., Bowtell, R., 2008. Simultaneous EEG source localisation and artifact rejection during concurrent fMRI by means of spatial filtering. Neuroimage 40, 1090–1104. https://doi.org/10.1016/j.neuroimage.2007.12.030

Brookes, M.J., Vrba, J., Mullinger, K.J., Geirsdóttir, G.B., Yan, W.X., Stevenson, C.M., Bowtell, R., Morris, P.G., 2009. Source localisation in concurrent EEG/fMRI: Applications at 7T. Neuroimage 45, 440–452. https://doi.org/10.1016/j.neuroimage.2008.10.047

Brown, P., Salenius, S., Rothwell, J.C., Hari, R., 1998. Cortical Correlate of the Piper Rhythm in Humans. J. Neurophysiol. 80, 2911–2917. https://doi.org/10.1152/jn.1998.80.6.2911

Buneo, C.A., Andersen, R.A., 2006. The posterior parietal cortex: Sensorimotor interface for the planning and online control of visually guided movements. Neuropsychologia 44, 2594–2606. https://doi.org/10.1016/j.neuropsychologia.2005.10.011

Buschman, T.J., Miller, E.K., 2007. Top-Down Versus Bottom-Up Control of Attention in the Prefrontal and Posterior Parietal Cortices. Science (80-.). 315, 1860–1862. https://doi.org/10.1126/science.1138071

Buzsáki, G., Anastassiou, C.A., Koch, C., 2012. The origin of extracellular fields and currents — EEG, ECoG, LFP and spikes. Nat. Rev. Neurosci. 13, 407–420. https://doi.org/10.1038/nrn3241

Buzsaki, G., Draguhn, A., 2004. Neuronal Oscillations in Cortical Networks. Science (80-.). 304, 1926–1929. https://doi.org/10.1126/science.1099745

Chandrasekaran, C., Canon, V., Dahmen, J.C., Kourtzi, Z., Welchman, A.E., 2007. Neural Correlates of Disparity-Defined Shape Discrimination in the Human Brain. J. Neurophysiol. 97, 1553–1565. https://doi.org/10.1152/jn.01074.2006

Cheyne, D., Bells, S., Ferrari, P., Gaetz, W., Bostan, A.C., 2008. Self-paced movements induce high-frequency gamma oscillations in primary motor cortex. Neuroimage 42, 332–342. https://doi.org/10.1016/j.neuroimage.2008.04.178

Cheyne, D., Ferrari, P., 2013. MEG studies of motor cortex gamma oscillations: evidence for a gamma “fingerprint” in the brain? Front. Hum. Neurosci. 7, 1–7. https://doi.org/10.3389/fnhum.2013.00575

Cheyne, D.O., 2013. MEG studies of sensorimotor rhythms: A review. Exp. Neurol. https://doi.org/10.1016/j.expneurol.2012.08.030

Cohen, M.X., 2017. Where Does EEG Come From and What Does It Mean? Trends Neurosci. 40, 208–218. https://doi.org/10.1016/j.tins.2017.02.004

Colgin, L.L., Denninger, T., Fyhn, M., Hafting, T., Bonnevie, T., Jensen, O., Moser, M.-B., Moser, E.I., 2009. Frequency of gamma oscillations routes flow of information in the hippocampus. Nature 462, 353–357. https://doi.org/10.1038/nature08573

Cottereau, B.R., McKee, S.P., Ales, J.M., Norcia, A.M., 2011. Disparity-Tuned Population Responses from Human Visual Cortex. J. Neurosci. 31, 954–965. https://doi.org/10.1523/JNEUROSCI.3795-10.2011

Crone, N.E., Miglioretti, D.L., Gordon, B., Lesser, R.P., 1998. Functional mapping of human sensorimotor cortex with electrocorticographic spectral analysis II. Event-related synchronization in the gamma band. Brain 121, 2301–2315. https://doi.org/10.1093/brain/121.12.2301

Culham, J.C., Danckert, S.L., De Souza, J.F.X., Gati, J.S., Menon, R.S., Goodale, M.A., 2003. Visually guided grasping produces fMRI activation in dorsal but not ventral stream brain areas. Exp. Brain Res. 153, 180–189. https://doi.org/10.1007/s00221-003-1591-5

Culham, J.C., Kanwisher, N.G., 2001. Neuroimaging of cognitive functions in human parietal cortex. Curr. Opin. Neurobiol. 11, 157–163. https://doi.org/10.1016/S0959-4388(00)00191-4

Cumming, B.G., 2002. An unexpected specialization for horizontal disparity in primate primary visual cortex. Nature 418, 633–636. https://doi.org/10.1038/nature00909

Cumming, B.G., DeAngelis, G.C., 2001. The Physiology of Stereopsis. Annu. Rev. Neurosci. 24, 203–238. https://doi.org/10.1146/annurev.neuro.24.1.203

Darvas, F., Scherer, R., Ojemann, J.G., Rao, R.P., Miller, K.J., Sorensen, L.B., 2010. High gamma mapping using EEG. Neuroimage 49, 930–938. https://doi.org/10.1016/j.neuroimage.2009.08.041

Delorme, A., Makeig, S., 2004. EEGLAB: an open source toolbox for analysis of single-trial EEG dynamics including independent component analysis. J. Neurosci. Methods 134, 9–21. https://doi.org/10.1016/j.jneumeth.2003.10.009

Delorme, A., Sejnowski, T., Makeig, S., 2007. Enhanced detection of artifacts in EEG data using higher-order statistics and independent component analysis. Neuroimage 34, 1443–1449. https://doi.org/10.1016/j.neuroimage.2006.11.004

Donner, T.H., Siegel, M., 2011. A framework for local cortical oscillation patterns. Trends Cogn. Sci. 15, 191–199. https://doi.org/10.1016/j.tics.2011.03.007

Durand, J.-B., Nelissen, K., Joly, O., Wardak, C., Todd, J.T., Norman, J.F., Janssen, P., Vanduffel, W., Orban, G.A., 2007. Anterior Regions of Monkey Parietal Cortex Process Visual 3D Shape. Neuron 55, 493–505. https://doi.org/10.1016/j.neuron.2007.06.040

Durand, J.-B., Peeters, R., Norman, J.F., Todd, J.T., Orban, G.A., 2009. Parietal regions processing visual 3D shape extracted from disparity. Neuroimage 46, 1114–1126. https://doi.org/10.1016/j.neuroimage.2009.03.023

Engel, A.K., Fries, P., Singer, W., 2001. Dynamic predictions: Oscillations and synchrony in top–down processing. Nat. Rev. Neurosci. 2, 704–716. https://doi.org/10.1038/35094565

Fell, J., Klaver, P., Lehnertz, K., Grunwald, T., Schaller, C., Elger, C.E., Fernández, G., 2001. Human memory formation is accompanied by rhinal–hippocampal coupling and decoupling. Nat. Neurosci. 4, 1259–1264. https://doi.org/10.1038/nn759

Freud, E., Macdonald, S.N., Chen, J., Quinlan, D.J., Goodale, M.A., Culham, J.C., 2018. Getting a grip on reality: Grasping movements directed to real objects and images rely on dissociable neural representations. Cortex 98, 34–48. https://doi.org/10.1016/j.cortex.2017.02.020

Freud, E., Plaut, D.C., Behrmann, M., 2016. ‘What’ Is Happening in the Dorsal Visual Pathway. Trends Cogn. Sci. 20, 773–784. https://doi.org/10.1016/j.tics.2016.08.003

Fries, P., 2015. Rhythms for Cognition: Communication through Coherence. Neuron 88, 220–235. https://doi.org/10.1016/j.neuron.2015.09.034

Fries, P., 2009. Neuronal Gamma-Band Synchronization as a Fundamental Process in Cortical Computation. Annu. Rev. Neurosci. 32, 209–224. https://doi.org/10.1146/annurev.neuro.051508.135603

Gaetz, W., MacDonald, M., Cheyne, D., Snead, O.C., 2010. Neuromagnetic imaging of movement-related cortical oscillations in children and adults: Age predicts post-movement beta rebound. Neuroimage 51, 792–807. https://doi.org/10.1016/j.neuroimage.2010.01.077

Gallivan, J.P., Culham, J.C., 2015. Neural coding within human brain areas involved in actions. Curr. Opin. Neurobiol. 33, 141–149. https://doi.org/10.1016/j.conb.2015.03.012

Gauthier, I., Hayward, W.G., Tarr, M.J., Anderson, A.W., Skudlarski, P., Gore, J.C., 2002. BOLD Activity during Mental Rotation and Viewpoint-Dependent Object Recognition. Neuron 34, 161–171. https://doi.org/10.1016/S0896-6273(02)00622-0

Georgieva, S., Peeters, R., Kolster, H., Todd, J.T., Orban, G.A., 2009. The Processing of Three-Dimensional Shape from Disparity in the Human Brain. J. Neurosci. 29, 727–742. https://doi.org/10.1523/JNEUROSCI.4753-08.2009

Georgieva, S.S., Todd, J.T., Peeters, R., Orban, G.A., 2008. The Extraction of 3D Shape from Texture and Shading in the Human Brain. Cereb. Cortex 18, 2416–2438. https://doi.org/10.1093/cercor/bhn002

Gilaie-Dotan, S., Ullman, S., Kushnir, T., Malach, R., 2002. Shape-selective stereo processing in human object-related visual areas. Hum. Brain Mapp. 15, 67–79. https://doi.org/10.1002/hbm.10008

Goncalves, N.R., Ban, H., Sanchez-Panchuelo, R.M., Francis, S.T., Schluppeck, D., Welchman, A.E., 2015. 7 Tesla fMRI Reveals Systematic Functional Organization for Binocular Disparity in Dorsal Visual Cortex. J. Neurosci. 35, 3056–3072. https://doi.org/10.1523/JNEUROSCI.3047-14.2015

Gonzalez, F., Perez, R., 1998. Neural mechanisms underlying stereoscopic vision. Prog. Neurobiol. 55, 191–224. https://doi.org/10.1016/S0301-0082(98)00012-4

Goodale, M.A., Milner, A.D., 2018. Two visual pathways – Where have they taken us and where will they lead in future? Cortex 98, 283–292. https://doi.org/10.1016/j.cortex.2017.12.002

Grefkes, C., Fink, G.R., 2005. The functional organization of the intraparietal sulcus in humans and monkeys. J. Anat. 207, 3–17. https://doi.org/10.1111/j.1469-7580.2005.00426.x

Gross, J., 2016. Let the Rhythm Guide You: Non-invasive Tracking of Cortical Communication Channels. Neuron 89, 244–247. https://doi.org/10.1016/j.neuron.2016.01.001

Haegens, S., Nacher, V., Luna, R., Romo, R., Jensen, O., 2011. α-Oscillations in the monkey sensorimotor network influence discrimination performance by rhythmical inhibition of neuronal spiking. Proc. Natl. Acad. Sci. 108, 19377–19382. https://doi.org/10.1073/pnas.1117190108

Hillebrand, A., Barnes, G.R., 2005. Beamformer Analysis of MEG Data. Int. Rev. Neurobiol. https://doi.org/10.1016/S0074-7742(05)68006-3

Hoogenboom, N., Schoffelen, J.-M., Oostenveld, R., Fries, P., 2010. Visually induced gamma-band activity predicts speed of change detection in humans. Neuroimage 51, 1162–1167. https://doi.org/10.1016/j.neuroimage.2010.03.041

Hoogenboom, N., Schoffelen, J.-M., Oostenveld, R., Parkes, L.M., Fries, P., 2006. Localizing human visual gamma-band activity in frequency, time and space. Neuroimage 29, 764–773. https://doi.org/10.1016/j.neuroimage.2005.08.043

Howard, M.W., Rizzuto, D.S., Caplan, J.B., Madsen, J.R., Lisman, J., Aschenbrenner-Scheibe, R., Schulze-Bonhage, A., Kahana, M.J., 2003. Gamma Oscillations Correlate with Working Memory Load in Humans. Cereb. Cortex 13, 1369–1374. https://doi.org/10.1093/cercor/bhg084

Hubbard, E.M., Piazza, M., Pinel, P., Dehaene, S., 2005. Interactions between number and space in parietal cortex. Nat. Rev. Neurosci. 6, 435–448. https://doi.org/10.1038/nrn1684

Jensen, O., Gips, B., Bergmann, T.O., Bonnefond, M., 2014. Temporal coding organized by coupled alpha and gamma oscillations prioritize visual processing. Trends Neurosci. 37, 357–369. https://doi.org/10.1016/j.tins.2014.04.001

Jensen, O., Mazaheri, A., 2010. Shaping Functional Architecture by Oscillatory Alpha Activity: Gating by Inhibition. Front. Hum. Neurosci. 4. https://doi.org/10.3389/fnhum.2010.00186

Joly, O., Vanduffel, W., Orban, G.A., 2009. The monkey ventral premotor cortex processes 3D shape from disparity. Neuroimage 47, 262–272. https://doi.org/10.1016/j.neuroimage.2009.04.043

Julesz, B., 1971. Foundations of Cyclopean Perception. Univ. Chicago Press, Chicago.

Jung, T.-P., Makeig, S., Humphries, C., Lee, T.-W., McKeown, M.J., Iragui, V., Sejnowski, T.J., 2000. Removing electroencephalographic artifacts by blind source separation. Psychophysiology 37, S0048577200980259. https://doi.org/10.1017/S0048577200980259

Kastner, S., Ungerleider, L.G., 2000. Mechanisms of Visual Attention in the Human Cortex. Annu. Rev. Neurosci. 23, 315–341. https://doi.org/10.1146/annurev.neuro.23.1.315

Kilavik, B.E., Zaepffel, M., Brovelli, A., MacKay, W.A., Riehle, A., 2013. The ups and downs of beta oscillations in sensorimotor cortex. Exp. Neurol. 245, 15–26. https://doi.org/10.1016/j.expneurol.2012.09.014

Klimesch, W., 1999. EEG alpha and theta oscillations reflect cognitive and memory performance: a review and analysis. Brain Res. Rev. 29, 169–195. https://doi.org/10.1016/S0165-0173(98)00056-3

Klimesch, W., Sauseng, P., Hanslmayr, S., 2007. EEG alpha oscillations: The inhibition–timing hypothesis. Brain Res. Rev. 53, 63–88. https://doi.org/10.1016/j.brainresrev.2006.06.003

Koenderink, J.J., 1998. Pictorial relief. Philos. Trans. R. Soc. A Math. Phys. Eng. Sci. 356, 1071–1086. https://doi.org/10.1098/rsta.1998.0211

Kourtzi, Z., Kanwisher, N., 2001. Representation of perceived object shape by the human lateral occipital complex. Science (80-.). 293, 1506–1509. https://doi.org/10.1126/science.1061133

Maris, E., 2012. Statistical testing in electrophysiological studies. Psychophysiology 49, 549–565. https://doi.org/10.1111/j.1469-8986.2011.01320.x

Maris, E., Oostenveld, R., 2007. Nonparametric statistical testing of EEG-and MEG-data. J. Neurosci. Methods 164, 177–190. https://doi.org/10.1016/j.jneumeth.2007.03.024

Mejias, J.F., Murray, J.D., Kennedy, H., Wang, X.-J., 2016. Feedforward and feedback frequency-dependent interactions in a large-scale laminar network of the primate cortex. Sci. Adv. 2, e1601335–e1601335. https://doi.org/10.1126/sciadv.1601335

Merriam, E.P., Colby, C.L., 2005. Active Vision in Parietal and Extrastriate Cortex. Neurosci. 11, 484–493. https://doi.org/10.1177/1073858405276871

Mesulam, M., Small, D., Vandenberg, R., Gitelman, D., Nobre, A., 2005. A heteromodal large-scale network for spatial attention, in: Itti, L., Rees, G., Tsotsos, J.K. (Eds.), Neurobiology of Attention. Elsevier, San Diego, pp. 29–34.

Michalareas, G., Vezoli, J., van Pelt, S., Schoffelen, J.-M., Kennedy, H., Fries, P., 2016. Alpha-Beta and Gamma Rhythms Subserve Feedback and Feedforward Influences among Human Visual Cortical Areas. Neuron 89, 384–397. https://doi.org/10.1016/j.neuron.2015.12.018

Michotte, A., 1991. The psychological enigma of perspective in outline pictures, in: Thines, G., Costall, A., Butterworth, G. (Eds.), Michotte’s Experimental Phenomenology of Perception. Erlbaum, Hillsdale, NJ, pp. 174–187.

Michotte, A., 1948. The Psychological Enigma of Perspective in Outline Pictures. Bull. la Cl. des Lettres l’Academie R. Belgique 34, 268–88.

Miller, K.J., Leuthardt, E.C., Schalk, G., Rao, R.P.N., Anderson, N.R., Moran, D.W., Miller, J.W., Ojemann, J.G., 2007. Spectral Changes in Cortical Surface Potentials during Motor Movement. J. Neurosci. 27, 2424–2432. https://doi.org/10.1523/JNEUROSCI.3886-06.2007

Minini, L., Parker, A.J., Bridge, H., 2010. Neural Modulation by Binocular Disparity Greatest in Human Dorsal Visual Stream. J. Neurophysiol. 104, 169–178. https://doi.org/10.1152/jn.00790.2009

Murray, S.O., Olshausen, B.A., Woods, D.L., 2003. Processing shape, motion and three-dimensional shape-from-motion in the human cortex. Cereb. Cortex 13, 508–516. https://doi.org/10.1093/cercor/13.5.508

Muthukumaraswamy, S.D., 2013. High-frequency brain activity and muscle artifacts in MEG/EEG: a review and recommendations. Front. Hum. Neurosci. 7, 1–11. https://doi.org/10.3389/fnhum.2013.00138

Muthukumaraswamy, S.D., 2010. Functional Properties of Human Primary Motor Cortex Gamma Oscillations. J. Neurophysiol. 104, 2873–2885. https://doi.org/10.1152/jn.00607.2010

Muthukumaraswamy, S.D., Singh, K.D., 2013. Visual gamma oscillations: The effects of stimulus type, visual field coverage and stimulus motion on MEG and EEG recordings. Neuroimage 69, 223–230. https://doi.org/10.1016/j.neuroimage.2012.12.038

Naganuma, T., Nose, I., Inoue, K., Takemoto, A., Katsuyama, N., Taira, M., 2005. Information processing of geometrical features of a surface based on binocular disparity cues: an fMRI study. Neurosci. Res. 51, 147–155. https://doi.org/10.1016/j.neures.2004.10.009

Neri, P., Bridge, H., Heeger, D.J., 2004. Stereoscopic Processing of Absolute and Relative Disparity in Human Visual Cortex. J. Neurophysiol. 92, 1880–1891. https://doi.org/10.1152/jn.01042.2003

Ni, J., Wunderle, T., Lewis, C.M., Desimone, R., Diester, I., Fries, P., 2016. Gamma-Rhythmic Gain Modulation. Neuron 92, 240–251. https://doi.org/10.1016/j.neuron.2016.09.003

Nichols, T.E., Holmes, A.P., 2002. Nonparametric permutation tests for functional neuroimaging: A primer with examples. Hum. Brain Mapp. 15, 1–25. https://doi.org/10.1002/hbm.1058

Ohzawa, I., DeAngelis, G., Freeman, R., 1990. Stereoscopic depth discrimination in the visual cortex: neurons ideally suited as disparity detectors. Science (80-.). 249, 1037–1041. https://doi.org/10.1126/science.2396096

Oostenveld, R., Fries, P., Maris, E., Schoffelen, J.M., 2011. FieldTrip: Open source software for advanced analysis of MEG, EEG, and invasive electrophysiological data. Comput Intell Neurosci 2011, 156869. https://doi.org/10.1155/2011/156869

Orban, G.A., 2011. The Extraction of 3D Shape in the Visual System of Human and Nonhuman Primates. Annu. Rev. Neurosci. 34, 361–388. https://doi.org/10.1146/annurev-neuro-061010-113819

Orban, G.A., Claeys, K., Nelissen, K., Smans, R., Sunaert, S., Todd, J.T., Wardak, C., Durand, J.-B., Vanduffel, W., 2006. Mapping the parietal cortex of human and non-human primates. Neuropsychologia 44, 2647–2667. https://doi.org/10.1016/j.neuropsychologia.2005.11.001

Orban, G.A., Sunaert, S., Todd, J.T., Van Hecke, P., Marchal, G., 1999. Human Cortical Regions Involved in Extracting Depth from Motion. Neuron 24, 929–940. https://doi.org/10.1016/S0896-6273(00)81040-5

Pantev, C., Makeig, S., Hoke, M., Galambos, R., Hampson, S., Gallen, C., 1991. Human auditory evoked gamma-band magnetic fields. Proc. Natl. Acad. Sci. 88, 8996–9000. https://doi.org/10.1073/pnas.88.20.8996

Parker, A.J., 2007. Binocular depth perception and the cerebral cortex. Nat. Rev. Neurosci. 8, 379–391. https://doi.org/10.1038/nrn2131

Pfurtscheller, G., Lopes da Silva, F.H., 1999. Event-related EEG/MEG synchronization and desynchronization: basic principles. Clin. Neurophysiol. 110, 1842–1857. https://doi.org/10.1016/S1388-2457(99)00141-8

Pfurtscheller, G., Stancák, A., Neuper, C., 1996. Event-related synchronization (ERS) in the alpha band — an electrophysiological correlate of cortical idling: A review. Int. J. Psychophysiol. 24, 39–46. https://doi.org/10.1016/S0167-8760(96)00066-9

Poggio, G.F., Motter, B.C., Squatrito, S., Trotter, Y., 1985. Responses of neurons in visual cortex (V1 and V2) of the alert macaque to dynamic random-dot stereograms. Vision Res. 25, 397–406. https://doi.org/10.1016/0042-6989(85)90065-3

Ponce, C.R., Born, R.T., 2008. Stereopsis. Curr. Biol. 18, R845–R850. https://doi.org/10.1016/j.cub.2008.07.006

Preston, T.J., Li, S., Kourtzi, Z., Welchman, A.E., 2008. Multivoxel Pattern Selectivity for Perceptually Relevant Binocular Disparities in the Human Brain. J. Neurosci. 28, 11315–11327. https://doi.org/10.1523/JNEUROSCI.2728-08.2008

Richter, C.G., Thompson, W.H., Bosman, C.A., Fries, P., 2017. Top-Down Beta Enhances Bottom-Up Gamma. J. Neurosci. 37, 6698–6711. https://doi.org/10.1523/JNEUROSCI.3771-16.2017

Robinson, S.E., Vrba, J., 1998. Functional neuroimaging by synthetic aperture magnetometry (SAM), in: Yoshimoto, T., Kotani, M., Kuriki, S., Karibe, H., Nakasato, N. (Eds.), Recent Advances in Biomagnetism. Tohoku Univ. Press, Sendai, Japan, pp. 302–305.

Rosenberg, A., Angelaki, D.E., 2014. Reliability-dependent contributions of visual orientation cues in parietal cortex. Proc. Natl. Acad. Sci. 111, 18043–18048. https://doi.org/10.1073/pnas.1421131111

Rosenberg, A., Cowan, N.J., Angelaki, D.E., 2013. The Visual Representation of 3D Object Orientation in Parietal Cortex. J. Neurosci. 33, 19352–19361. https://doi.org/10.1523/JNEUROSCI.3174-13.2013

Sakata, H., Tsutsui, K.-I., Taira, M., 2005. Toward an understanding of the neural processing for 3D shape perception. Neuropsychologia 43, 151–161. https://doi.org/10.1016/j.neuropsychologia.2004.11.003

Schadow, J., Lenz, D., Dettler, N., Fründ, I., Herrmann, C.S., 2009. Early gamma-band responses reflect anticipatory top-down modulation in the auditory cortex. Neuroimage 47, 651–658. https://doi.org/10.1016/j.neuroimage.2009.04.074

Scheeringa, R., Fries, P., Petersson, K.-M., Oostenveld, R., Grothe, I., Norris, D.G., Hagoort, P., Bastiaansen, M.C.M., 2011. Neuronal Dynamics Underlying High- and Low-Frequency EEG Oscillations Contribute Independently to the Human BOLD Signal. Neuron 69, 572–583. https://doi.org/10.1016/j.neuron.2010.11.044

Scheeringa, R., Koopmans, P.J., van Mourik, T., Jensen, O., Norris, D.G., 2016. The relationship between oscillatory EEG activity and the laminar-specific BOLD signal. Proc. Natl. Acad. Sci. 113, 6761–6766. https://doi.org/10.1073/pnas.1522577113

Schlosberg, H., 1941. Stereoscopic Depth from Single Pictures. Am. J. Psychol. 54, 601–605. https://doi.org/10.2307/1417214

Schoffelen, J., Oostenveld, R., Fries, P., 2005. Neuronal coherence as a mechanism of effective corticospinal interaction. Science 308, 111–113. https://doi.org/10.1126/science.1107027

Sejnowski, T.J., Churchland, P.S., Movshon, J.A., 2014. Putting big data to good use in neuroscience. Nat. Neurosci. 17, 1440–1441. https://doi.org/10.1038/nn.3839

Shikata, E., Hamzei, F., Glauche, V., Knab, R., Dettmers, C., Weiller, C., Büchel, C., 2001. Surface Orientation Discrimination Activates Caudal and Anterior Intraparietal Sulcus in Humans: An Event-Related fMRI Study. J. Neurophysiol. 85, 1309–1314. https://doi.org/10.1152/jn.2001.85.3.1309

Shikata, E., Hamzei, F., Glauche, V., Koch, M., Weiller, C., Binkofski, F., Büchel, C., 2003. Functional properties and interaction of the anterior and posterior intraparietal areas in humans. Eur. J. Neurosci. 17, 1105–1110. https://doi.org/10.1046/j.1460-9568.2003.02540.x

Siegel, M., Donner, T.H., Engel, A.K., 2012. Spectral fingerprints of large-scale neuronal interactions. Nat. Rev. Neurosci. 13, 121–134. https://doi.org/10.1038/nrn3137

Singer, W., Gray, C.M., 1995. Visual Feature Integration and the Temporal Correlation Hypothesis. Annu. Rev. Neurosci. 18, 555–586. https://doi.org/10.1146/annurev.neuro.18.1.555

Snow, J.C., Pettypiece, C.E., McAdam, T.D., McLean, A.D., Stroman, P.W., Goodale, M.A., Culham, J.C., 2011. Bringing the real world into the fMRI scanner: Repetition effects for pictures versus real objects. Sci. Rep. 1, 130. https://doi.org/10.1038/srep00130

Spaak, E., Bonnefond, M., Maier, A., Leopold, D.A., Jensen, O., 2012. Layer-Specific Entrainment of Gamma-Band Neural Activity by the Alpha Rhythm in Monkey Visual Cortex. Curr. Biol. 22, 2313–2318. https://doi.org/10.1016/j.cub.2012.10.020

Spang, K., Gillam, B., Fahle, M., 2012. Electrophysiological Correlates of Binocular Stereo Depth without Binocular Disparities. PLoS One 7, e40562. https://doi.org/10.1371/journal.pone.0040562

Taira, M., Nose, I., Inoue, K., Tsutsui, K., 2001. Cortical Areas Related to Attention to 3D Surface Structures Based on Shading: An fMRI Study. Neuroimage 14, 959–966. https://doi.org/10.1006/nimg.2001.0895

Taira, M., Tsutsui, K.-I., Jiang, M., Yara, K., Sakata, H., 2000. Parietal Neurons Represent Surface Orientation From the Gradient of Binocular Disparity. J. Neurophysiol. 83, 3140–3146. https://doi.org/10.1152/jn.2000.83.5.3140

Thut, G., Miniussi, C., 2009. New insights into rhythmic brain activity from TMS–EEG studies. Trends Cogn. Sci. 13, 182–189. https://doi.org/10.1016/j.tics.2009.01.004

Tsao, D.Y., Vanduffel, W., Sasaki, Y., Fize, D., Knutsen, T.A., Mandeville, J.B., Wald, L.L., Dale, A.M., Rosen, B.R., Van Essen, D.C., Livingstone, M.S., Orban, G.A., Tootell, R.B.H., 2003. Stereopsis Activates V3A and Caudal Intraparietal Areas in Macaques and Humans. Neuron 39, 555–568. https://doi.org/10.1016/S0896-6273(03)00459-8

Tsutsui, K.I., Sakata, H., Naganuma, T., Taira, M., 2002. Neural correlates for perception of 3D surface orientation from texture gradient. Science (80-.). 298, 409–412. https://doi.org/10.1126/science.1074128

Tunik, E., Rice, N.J., Hamilton, A., Grafton, S.T., 2007. Beyond grasping: Representation of action in human anterior intraparietal sulcus. Neuroimage 36, T77–T86. https://doi.org/10.1016/j.neuroimage.2007.03.026

Uji, M., Wilson, R., Francis, S.T., Mullinger, K.J., Mayhew, S.D., 2018. Exploring the advantages of multiband fMRI with simultaneous EEG to investigate coupling between gamma frequency neural activity and the BOLD response in humans. Hum. Brain Mapp. 39, 1673–1687. https://doi.org/10.1002/hbm.23943

Van Dromme, I.C., Premereur, E., Verhoef, B.-E., Vanduffel, W., Janssen, P., 2016. Posterior Parietal Cortex Drives Inferotemporal Activations During Three-Dimensional Object Vision. PLoS Biol. 14, e1002445. https://doi.org/10.1371/journal.pbio.1002445

Van Dromme, I.C.L., Vanduffel, W., Janssen, P., 2015. The relation between functional magnetic resonance imaging activations and single-cell selectivity in the macaque intraparietal sulcus. Neuroimage 113, 86–100. https://doi.org/10.1016/j.neuroimage.2015.03.023

van Drongelen, W., Yuchtman, M., Van Veen, B.D., van Huffelen, A.C., 1996. A spatial filtering technique to detect and localize multiple sources in the brain. Brain Topogr. 9, 39–49. https://doi.org/10.1007/BF01191641

van Kerkoerle, T., Self, M.W., Dagnino, B., Gariel-Mathis, M.-A., Poort, J., van der Togt, C., Roelfsema, P.R., 2014. Alpha and gamma oscillations characterize feedback and feedforward processing in monkey visual cortex. Proc. Natl. Acad. Sci. 111, 14332–14341. https://doi.org/10.1073/pnas.1402773111

van Veen, B.D., van Drongelen, W., Yuchtman, M., Suzuki, A., 1997. Localization of brain electrical activity via linearly constrained minimum variance spatial filtering. Biomedical 44, 867–880. https://doi.org/10.1109/10.623056

Vanduffel, W., Fize, D., Peuskens, H., Denys, K., Sunaert, S., Todd, J.T., Orban, G.A., 2002. Extracting 3D from motion: Differences in human and monkey intraparietal cortex. Science (80-.). 298, 413–415. https://doi.org/10.1126/science.1073574

Verhoef, B.-E., Vogels, R., Janssen, P., 2011. Synchronization between the end stages of the dorsal and the ventral visual stream. J. Neurophysiol. 105, 2030–2042. https://doi.org/10.1152/jn.00924.2010

Verhoef, B.E., Bohon, K.S., Conway, B.R., 2015. Functional Architecture for Disparity in Macaque Inferior Temporal Cortex and Its Relationship to the Architecture for Faces, Color, Scenes, and Visual Field. J. Neurosci. 35, 6952–6968. https://doi.org/10.1523/JNEUROSCI.5079-14.2015

Vishwanath, D., 2014. Toward a new theory of stereopsis. Psychol. Rev. 121, 151–178. https://doi.org/10.1037/a0035233

Vishwanath, D., Hibbard, P.B., 2013. Seeing in 3-D With Just One Eye: Stereopsis in the absence of binocular disparities. Psychol. Sci. 24, 1673–1685. https://doi.org/10.1177/0956797613477867

Volcic, R., Vishwanath, D., Domini, F., 2014. Reaching into Pictorial Spaces, in: Rogowitz, B.E., Pappas, T.N., de Ridder, H. (Eds.), Human Vision and Electronic Imaging XIX (Vol. 9014). International Society for Optics and Photonics. p. 901413. https://doi.org/10.1117/12.2045458

Wade, N.J., Ono, H., Lillakas, L., 2001. Leonardo da Vinci’s Struggles with Representations of Reality. Leonardo 34, 231–235. https://doi.org/10.1162/002409401750286994

Wagner, A.D., Shannon, B.J., Kahn, I., Buckner, R.L., 2005. Parietal lobe contributions to episodic memory retrieval. Trends Cogn. Sci. 9, 445–453. https://doi.org/10.1016/j.tics.2005.07.001

Wan, X., Sekiguchi, A., Yokoyama, S., Riera, J., Kawashima, R., 2008. Electromagnetic source imaging: Backus–Gilbert resolution spread function-constrained and functional MRI-guided spatial filtering. Hum. Brain Mapp. 29, 627–643. https://doi.org/10.1002/hbm.20424

Wang, X.-J., 2010. Neurophysiological and Computational Principles of Cortical Rhythms in Cognition. Physiol. Rev. 90, 1195–1268. https://doi.org/10.1152/physrev.00035.2008

Watt, S.J., Bradshaw, M.F., 2003. The visual control of reaching and grasping: Binocular disparity and motion parallax. J. Exp. Psychol. Hum. Percept. Perform. 29, 404–415. https://doi.org/10.1037/0096-1523.29.2.404

Welchman, A.E., 2016. The Human Brain in Depth: How We See in 3D. Annu. Rev. Vis. Sci. 2, 345–376. https://doi.org/10.1146/annurev-vision-111815-114605

Welchman, A.E., Kourtzi, Z., 2013. Linking brain imaging signals to visual perception. Vis. Neurosci. 30, 229–241. https://doi.org/10.1017/S0952523813000436

Westheimer, G., 2011. Three-dimensional displays and stereo vision. Proc. R. Soc. B Biol. Sci. 278, 2241–2248. https://doi.org/10.1098/rspb.2010.2777

Wheatstone, C., 1838. Contributions to the Physiology of Vision. Part the First. On Some Remarkable, and Hitherto Unobserved, Phenomena of Binocular Vision. Philos. Trans. R. Soc. London 128, 371–394. https://doi.org/10.1098/rstl.1838.0019

Womelsdorf, T., Fries, P., Mitra, P.P., Desimone, R., 2006. Gamma-band synchronization in visual cortex predicts speed of change detection. Nature 439, 733–736. https://doi.org/10.1038/nature04258

Yantis, S., Serences, J.T., 2003. Cortical mechanisms of space-based and object-based attentional control. Curr. Opin. Neurobiol. 13, 187–193. https://doi.org/10.1016/S0959-4388(03)00033-3

